# Identification of the negamycin split biosynthetic gene cluster in *Kitasatospora purpeofusca* ATCC21470

**DOI:** 10.1101/2025.07.31.667836

**Authors:** Alena Strüder, Saad Touzi, Mayarasmi Long, Erik Mingyar, Carolin Schwitalla, Anne Berscheid, Sophie Bourcier, Heike Brötz-Oesterhelt, Emmanuelle Darbon, Paul Frankenreiter, Mirita Franz-Wachtel, Christian Geibel, Theresa Harbig, Chambers C. Hughes, Boris Macek, Kay Nieselt, Helena Sales-Ortells, David Touboul, Sylvie Lautru, Evi Stegmann

**Author notes:** equal contribution (Alena Strüder, Saad Touzi). Senior and Corresponding authors: ES +SL.

## Abstract

Negamycin is a ribosome-targeting antibiotic with activity against Gram-positive and Gram-negative bacteria including ESKAPE pathogens. Furthermore, it promotes premature stop codon readthrough. Its therapeutic potential is limited by low natural production and synthetic complexity. To enable scalable biosynthesis, we identified and characterized its genetic basis in *Kitasatospora purpeofusca* ATCC 21470. Two distant chromosomal regions, *neg1* and *neg2*, were found to be essential. Deletion of *neg1*, involved in nitrite provision for N-N bond formation, reduced production to ∼10%, while deletion of *neg2*, which directs β-lysine generation and scaffold assembly, abolished it completely. Isotope-labeling experiments confirmed nitrite incorporation. Transcriptomic and proteomic analyses further supported the involvement of both regions. The heterologous expression of *neg1* along with the *neg2* region in *Streptomyces albidoflavus* reconstituted negamycin biosynthesis, confirming the unusual involvement of two distant gene clusters in the biosynthesis, and provides a foundation for biotechnological production and further development of this promising antibiotic.

**SIGNIFICANCE:** The rapid rise of antimicrobial resistance (AMR), particularly among Gram-negative ESKAPE pathogens, represents one of the most urgent global health threats. Despite this, the discovery and development of new antibiotics have stagnated. Addressing this challenge requires the exploration of natural products with novel mechanisms of action, alongside the development of scalable production strategies.

Negamycin has emerged as a compelling candidate in this regard, characterized by an unusual mechanism of action and therapeutic potential extending beyond traditional antibacterial use. However, its development has been constrained by low production yields in the native producer. In this study, we identify and characterize the biosynthetic genes responsible for negamycin production, providing a foundation for pathway engineering, yield optimization, and the rational design of new analogs.

## INTRODUCTION

The ongoing emergence of multidrug-resistant pathogens poses a critical threat to global public health and underscores the urgent need for new antibiotics with novel mechanisms of action. Negamycin, a naturally occurring hydrazide compound first isolated in 1970 from the culture supernatant of *Streptomyces purpeofuscus* (now *Kitasatospora purpeofusca*), exhibits a broad spectrum of antibacterial activity against both Gram-negative and Gram-positive bacteria, including clinically relevant members of the ESKAPE group such as *Klebsiella pneumoniae*, *Enterobacter spp.*, and *Pseudomonas aeruginosa*^1,2^. Interestingly, negamycin shows enhanced potency against Gram-negative bacteria^3^, a rare and highly desirable trait among current antibiotics. In addition to its antibacterial properties, negamycin has attracted considerable interest for its ability to induce translational readthrough of premature termination codons (PTCs), a mechanism with therapeutic potential for treating a wide range of genetic disorders caused by nonsense mutations, including Duchenne muscular dystrophy (DMD), cystic fibrosis (CF), hemophilia A, and certain cancers^4–6^.

Negamycin functions by a unique mechanism of action; it binds to the helix 34 of the 16S rRNA of the bacterial ribosome at a site distinct from those targeted by aminoglycosides or tetracyclines, establishing interactions with both the rRNA and aminoacyl-tRNA^7^. This binding increases the residence time of near-cognate tRNAs, leading to miscoding and bactericidal activity. Notably, this novel binding mode helps circumvent existing resistance mechanisms that impair other ribosome-targeting antibiotics. Structure-activity relationship (SAR) studies have revealed that while analogs with enhanced readthrough activity are relatively easy to generate^8–11^, improving antibacterial efficacy has proven more challenging. For instance, N6-(3-aminopropyl)-negamycin is one of the few derivatives that display improved antimicrobial properties^10^. In contrast, natural analogs such as 3-epi-deoxynegamycin and leucyl-3-epi-deoxynegamycin show significantly higher readthrough activity, yet lack antibacterial activities, making them promising candidates for PTC suppression therapies^12^.

Despite its therapeutic potential, both as an antibiotic and as a readthrough agent, large-scale application of negamycin and its analogs remains limited due to the low productivity of native actinobacterial producers and the complexity of chemical synthesis^13^. The compound’s structure is highly unusual, featuring a rare N-N bond, which contributes to both its biological activity and its synthetic intractability.

Over the past few years, significant progress has been made in elucidating the biosynthesis of N-N bond-containing natural products^14^. Three main biosynthetic strategies have been identified. One involves the production of nitrous acid (HNO₂) via the aspartate/nitrosuccinate (ANS) pathway, as exemplified in cremeomycin biosynthesis, where enzymes CreE and CreD generate HNO₂ that is subsequently incorporated into the metabolite scaffold by the ligase CreM^15^. Another route entails the formation of N-hydroxylated amino acids through the activity of flavin-dependent monooxygenases, such as in the biosynthesis of piperazic acid, a key building block in kutzneride and triacsin biosynthesis^16,17^. A third strategy involves the biosynthesis of hydrazinoacetic acid (HAA), catalyzed by enzymes such as Spb38–Spb40 in the s56-p1 pathway, where the final N-N bond is formed via an ATP-dependent condensation of amino acid substrates, followed by rearrangement facilitated by a zinc-binding cupin domain^18,19^.

Given its unusual structure and promising bioactivity, a deeper understanding of negamycin biosynthesis is crucial. Elucidating the underlying biosynthetic pathway would not only facilitate strain engineering for increased production yields but also provide a foundation for the generation of novel derivatives with enhanced antibacterial or readthrough properties. During the final stages of our work, an independent study was published^20^, presenting the identification of the negamycin biosynthetic genes and key insights into its assembly. Our findings are in strong agreement with those reported and further corroborate them through additional genetic, transcriptional experiments and proteomic analysis. This parallel discovery highlights the robustness of the current model and underscores the importance of understanding N-N bond biosynthesis in natural products.

## RESULTS

### Genome mining reveals two candidate regions for negamycin biosynthesis

To elucidate the genetic basis for negamycin biosynthesis, we sequenced the genome of the producing strain *Kitasatospora purpeofusca* ATCC 21470 using a PacBio sequencing platform. De novo assembly with HGAP3 (Hierarchical Genome Assembly Process) yielded two contigs, which were subsequently scaffolded using the genome of *K. purpeofusca* NRBC 14457 as a reference. To validate the contig junctions, the corresponding genomic region was PCR-amplified and confirmed by Sanger sequencing. Genome annotation was performed using the RAST annotation server^21, 22^.

To identify potential biosynthetic loci, we subjected the annotated genome to antiSMASH^23^, a tool that predicts biosynthetic gene clusters (BGCs) based on conserved biosynthetic signatures. This analysis revealed 39 putative BGCs. However, manual inspection failed to identify a candidate that could plausibly encode the biosynthesis of negamycin.

Prompted by the distinctive structural features of negamycin, specifically a glycine and β-lysine moiety connected via a hydrazide bond (Figure 1), we pursued a targeted genome mining approach to identify genes encoding enzymes capable of catalyzing N-N bond formation.

**Figure 1:**
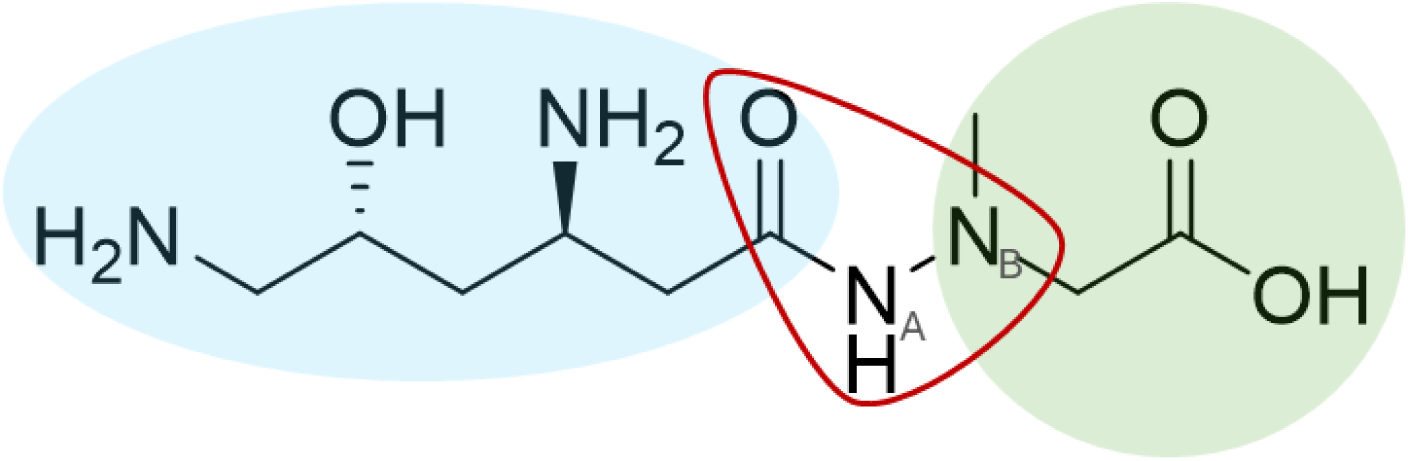
Structure of negamycin. Negamycin is a pseudopeptidic hydrazide compound containing a δ-OH-β-lysine (blue) and an N-methyl-glycine (green) connected via a specific N-N-bond forming a hydrazide moiety (red). A: N atom derived from nitrous acid; B: N atom derived from glycine.

To this end, the *K. purpeofusca* genome was screened for genes homologous to those encoding known N-N bond-forming enzymes. This search yielded two candidate loci. The first region, located between positions 4,726,499 and 4,732,858 bp, contains genes encoding enzymes with high sequence similarity to CreM, CreE, and CreD (KPATCC21470_4344, _4346, _4347) (Figure 2; Table S1 and S2), involved, in cremeomycin biosynthesis, in the generation of the nitrous acid precursor by the aspartate-nitrosuccinate (ANS) pathway (CreE and CreD) and in N-N bond formation (CreM). The second region (9,557,759 to 9,564,420 bp, Figure S1), was identified when searching for genes encoding homologs of Spb38, Spb39, and Spb40 (Table S1), responsible for the biosynthesis of the hydrazinoacetic acid precursor in S56-p1 biosynthesis. This region encodes homologs of Spb38 (KPATCC21470_8504), Spb39 (KPATCC21470_8500), and Spb40 (KPATCC21470_8499).

**Figure 2:**
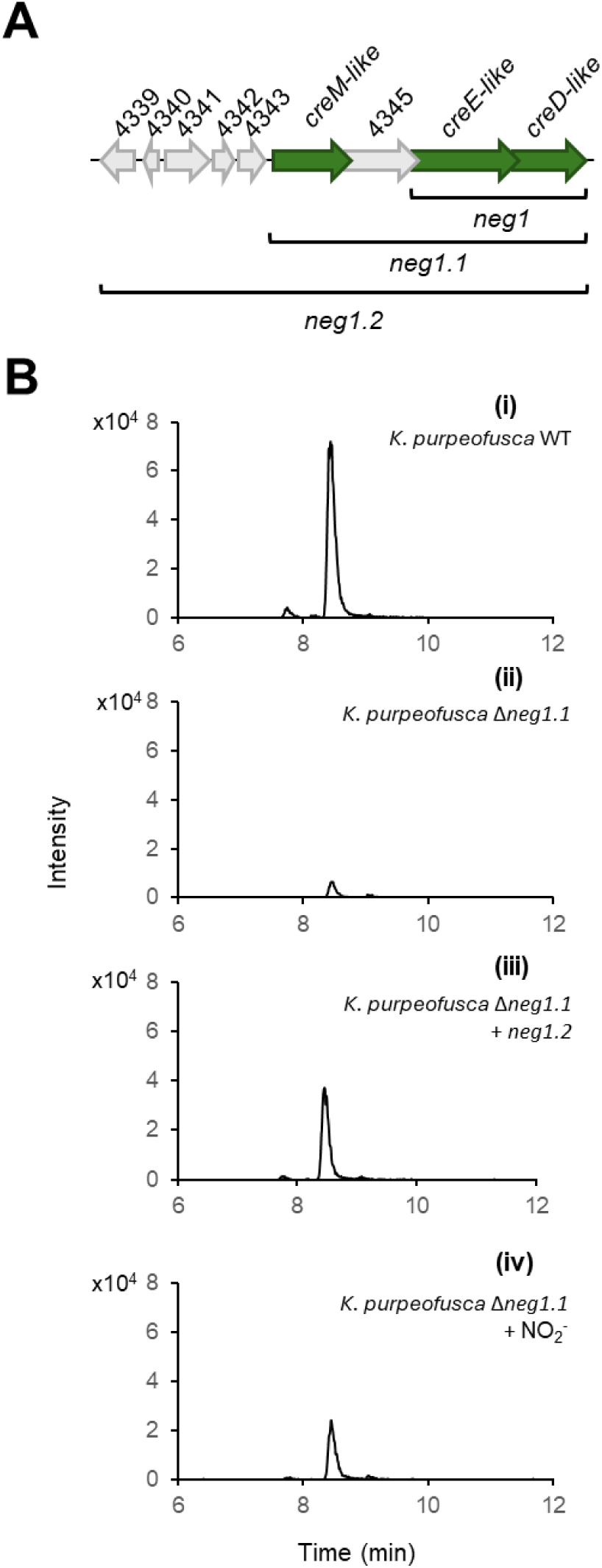
Genomic region 1, putatively involved in negamycin biosynthesis (A), and LC-ESI-HRMS analysis of negamycin production. **(B)** A: Genes highlighted in green encode homologs of enzymes involved in nitrous acid generation (*creE-*like and *creD-like,* encoding an N-monooxygenase and an aspartase, respectively) and N-N bond formation (*creM-like,* encoding a ligase) known from cremeomycin biosynthesis in *Streptomyces cremeus*. KPATCC21470_4345 (encoding a FAD-dependent monooxygenase) is not present in the cremeomycin biosynthetic gene cluster. *neg1.1* corresponds to the region deleted in the *K. purpeofusca* Δ*neg1.1* mutant. *neg1.2* corresponds to the region used for the genetic complementation of *K. purpeofusca* Δ*neg1.1*. *neg1* encompasses the *creE* and *creD-*like genes involved in negamycin biosynthesis and responsible for nitrite supply. B: Extracted ion chromatogram (*m/z* 471.2238 [M+H]^+^) of partially purified and FMOC-derivatized culture supernatants (see STAR Methods) of (i) *K. purpeofusca* ATCC 21470; (ii) *K. purpeofusca* Δ*neg1.1* mutant; (ii) Genetically complemented *K. purpeofusca* Δ*neg1.1* mutant; and (iv) *K. purpeofusca* Δ*neg1.1* supplemented with nitrite. See also Figure S2 and Table S2.

### Negamycin biosynthesis involves region 1 but not region 2

To evaluate the involvement of the region 1 (including *cre*-like genes) and/or region 2 (*spb*-like genes) in negamycin biosynthesis, we first performed targeted deletions. Deletion of the *spb-like* genes (KPATCC21470_8499-8504) (Figure S1) via classical two-step homologous recombination had no effect on negamycin production (data not shown), indicating that region 2 does not contribute to the biosynthesis of negamycin. In contrast, deletion of four genes from region 1 (KPATCC21470_4344-4347, designated as *neg1.1* region, Figure 2,A) using the same strategy led to a near-complete loss of negamycin production in the resulting mutant *K. purpeofusca* Δ*neg1.1*, with residual levels reduced to approximately 9% based on peak area (Figure 2, B (i), (ii), and Figure S2). These findings implicate region 1 in negamycin biosynthesis.

To verify that the reduced negamycin production was specifically due to the KPATCC21470_4344-4347 deletion, we genetically complemented the *K. purpeofusca* Δ*neg1.1* mutant with plasmid pRM4_*neg1.2*. This construct contains not only the deleted genes but also the four additional genes KPATCC21470_4339, _4340, _4342, _4343, upstream of the *neg1.1,* which may encode accessory functions (Figure 2, A). Complementation restored negamycin biosynthesis, resulting in a 30% increase in peak area relative to the *K. purpeofusca Δneg1.1* mutant (Figure 2, B(iii)). These results confirmed that the *neg1.1* region is essential for negamycin biosynthesis.

### Nitrite is a precursor in negamycin biosynthesis

The involvement of the *neg1.1* region, particularly the homologs of *creE* and *creD*, suggested that precursor supply in negamycin biosynthesis proceeds via the L-aspartate-nitrosuccinate (ANS) pathway, implicating nitrite as a biosynthetic precursor. To test this hypothesis, we supplemented cultures of *K. purpeofusca* with **^1^**^5^N-labeled nitrite (^15^NO_2_ ^−^). After partial purification by solid phase extraction and FMOC derivatization, we analyzed the culture supernatants by LC-ESI-HRMS. The mass spectrum showed a clear isotopic shift, with a 77% enrichment in the M+1 isotope (Figure S3), providing strong evidence for the incorporation of the ^15^N atom from ^15^NO_2_ ^−^ into negamycin and supporting nitrite as a biosynthetic precursor.

We further examined whether nitrite supplementation could restore negamycin production in the *K. purpeofusca* Δ*neg1.1*. Indeed, addition of nitrite to the culture medium led to a partial rescue of negamycin biosynthesis, with production levels reaching ∼50% of wild-type *K. purpeofusca* (in peak area, Figure 2, B (iv)). The residual negamycin production observed in *K. purpeofusca* Δ*neg1.1* cultures, along with its partial restoration upon nitrite supplementation, suggested that trace amounts of nitrite inherently present in the production medium might account for the low-level biosynthesis. To test this, we analyzed individual medium components using the Griess assay and detected low concentrations of nitrite in both soy flour and yeast extract (Figure S4), providing a plausible explanation for the residual negamycin detected in the mutant culture supernatants.

This finding not only confirms the role of nitrite as a biosynthetic precursor but also suggests that among the four deleted genes, only those encoding homologs of CreE and CreD are directly required for precursor formation.

### Only the *creE*- and *creD*-like genes in *neg1.1* are expressed during negamycin production

Since in *neg1.1* only KPATCC21470_4346 and KPATCC21470_4347 appeared to be involved in negamycin biosynthesis, we considered the possibility that additional biosynthetic routes might contribute to negamycin production. To explore this, we conducted a transcriptomic analysis of *K. purpeofusca* wild type strain after 96 hours of cultivation under production conditions. Total RNA was extracted, and genome-wide transcriptional profile was analyzed.

Within the *neg1.1* locus, only the *creE*-like gene (KPATCC21470_4346) showed detectable transcriptional activity at this time point (Figure S5). Neither *creD*-like nor *creM*-like genes (encoding an ATP-dependent ligase) were expressed. Moreover, none of the *spb*-like genes (KPUR_ATCC21470_08504, 08500, 08499) were transcribed. These findings are consistent with the deletion experiments, further supporting that the region 2 is not involved in negamycin biosynthesis.

To validate and temporally resolve these observations, we performed RT-PCR using cDNA from *K. purpeofusca* wild type cultures harvested at 24, 36 and 48 hours. This analysis confirmed constant expression of *creE*-like and of *creD*-like genes (KPATCC21470_4346, 4347) at all time points (Figure S6). In contrast, *creM*-like and all tested *spb* homologs remained transcriptionally silent throughout.

Together, these results demonstrate that negamycin biosynthesis under standard laboratory conditions relies on the expression of *creE-*like and *creD*-like within the *neg1.1* locus. The region encompassing these two genes was therefore designated *neg1,* to reflect its functional contribution to the negamycin biosynthetic pathway.

### L-lysine and glycine are negamycin precursors

To determine the origin of the other structural components of negamycin, the β-hydroxy-lysine moiety and the N-methylglycine moiety, likely deriving from L-lysine and glycine, respectively, we investigated their biosynthetic sources. Cultures of *K. purpeofusca* were supplemented with ¹³C₆ - labeled L-lysine, ¹³C₂-labeled glycine, or ¹⁵N-labeled glycine (see STAR Methods). Following solid-phase extraction and FMOC derivatization, culture supernatants were analyzed by LC-ESI-HRMS. In ¹³C₆ -lysine-fed cultures, we observed the incorporation of all ^13^C from ¹³C₆ -lysine as demonstrated by the M+6 isotopic peak (61% enrichment, Figure S7), supporting the hypothesis that L-lysine is a precursor of negamycin. ¹³C₂- and ¹⁵N-glycine-fed cultures showed an enrichment of the M+2 (9% enrichment) and M+1 (18% enrichment, Figure S7) isotopic peaks respectively, suggesting that glycine is incorporated intact into the negamycin scaffold and that the N atom derived from nitrite is N_A_ (Figure 1).

### The β-lysine precursor is synthesized by a distinct gene region outside the *neg1* locus

As L-lysine was confirmed as a precursor of negamycin, and since β-lysine is known to arise from L-lysine via the enzymatic action of lysine 2,3-aminomutases^24^, we sought to identify the specific gene(s) responsible for this transformation in *K. purpeofusca* genome.

Genome analysis revealed three candidate genes encoding putative lysine 2,3-aminomutases: *kamA1* (*KPATCC21470_2073*), *kamA2* (*KPATCC21470_8074*), and *kamA3* (*KPATCC21470_8180*). RT-PCR analysis after growing for 24 h, 36 h and 48 h under negamycin producing conditions showed that only *kamA2* was transcriptionally active, while *kamA1* and *kamA3* remained silent (Figure S6). These findings suggested that *kamA2* was solely responsible for β-lysine synthesis in the negamycin pathway.

To further support the functional relevance of *kamA2*, we performed comparative proteomic analysis of the *K. purpeofusca* wild type strain and the *K. purpeofusca* Δ*neg1.1* mutant. Both strains were cultivated under production conditions, and samples were harvested after 72 hours. We systematically compared the proteomes of *K. purpeofusca* wild type and *K. purpeofusca* Δ*neg1.1*. Notably, KamA2 and four proteins encoded in the vicinity of *kamA2* (*KPATCC21470_8071*–*8075*) were significantly more abundant in the wild type strain (Table S3). Encouraged by these results, we examined the expression of the corresponding genes in our transcriptomic dataset. We found that all five genes are expressed under negamycin-producing conditions and are part of an 11 kb transcriptionally active region spanning KPATCC21470_8065 to KPATCC21470_8075 (Figure S5, C). In addition to *kamA2,* encoding the lysine 2,3-aminomutase, this region also contains KPATCC21470_8066, encoding a protein homologous to KinJ (29.6% identity), an enzyme involved in N-N bond formation in kinamycin biosynthesis. This suggests that these genes may constitute a second, functionally distinct biosynthetic locus required for negamycin production, contributing to both the β-lysine precursor and potentially an N-N bond-forming activity. If this were the case, the region would likely be conserved in other negamycin-producing bacteria, as BGCs are often disseminated via horizontal gene transfer.

To test this, we used the Cluseek 1.4.26 program (https://lab111.mbu.cas.cz/cluseek/) to assess whether these 11 genes are colocalized in other bacterial genomes. All genes except KPATCC21470_8065 were conserved across seven species of Actinomycetota, forming a genomic island (Figure 3). These findings support the hypothesis that KPATCC21470_8066 and KPATCC21470_8075 define the boundaries of a second segment of the negamycin BGC, which we designated as *neg2* (Figure 4, A).

**Figure 3:**
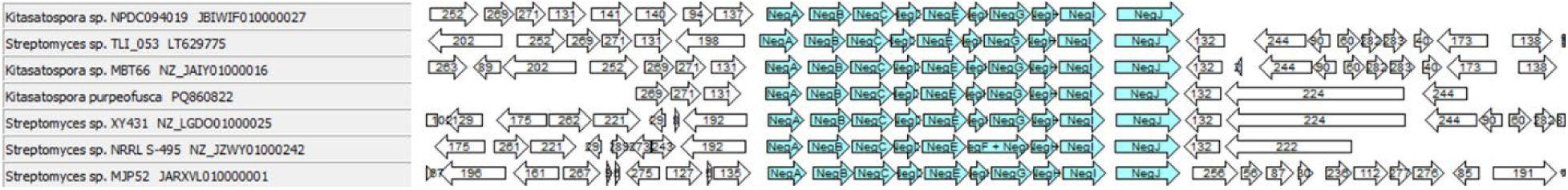
Genetic context of the *neg2* region (in blue) in seven actinomycetota. The figure was created using CluSeek 1.4.26 (https://lab111.mbu.cas.cz/cluseek/)

**Figure 4:**
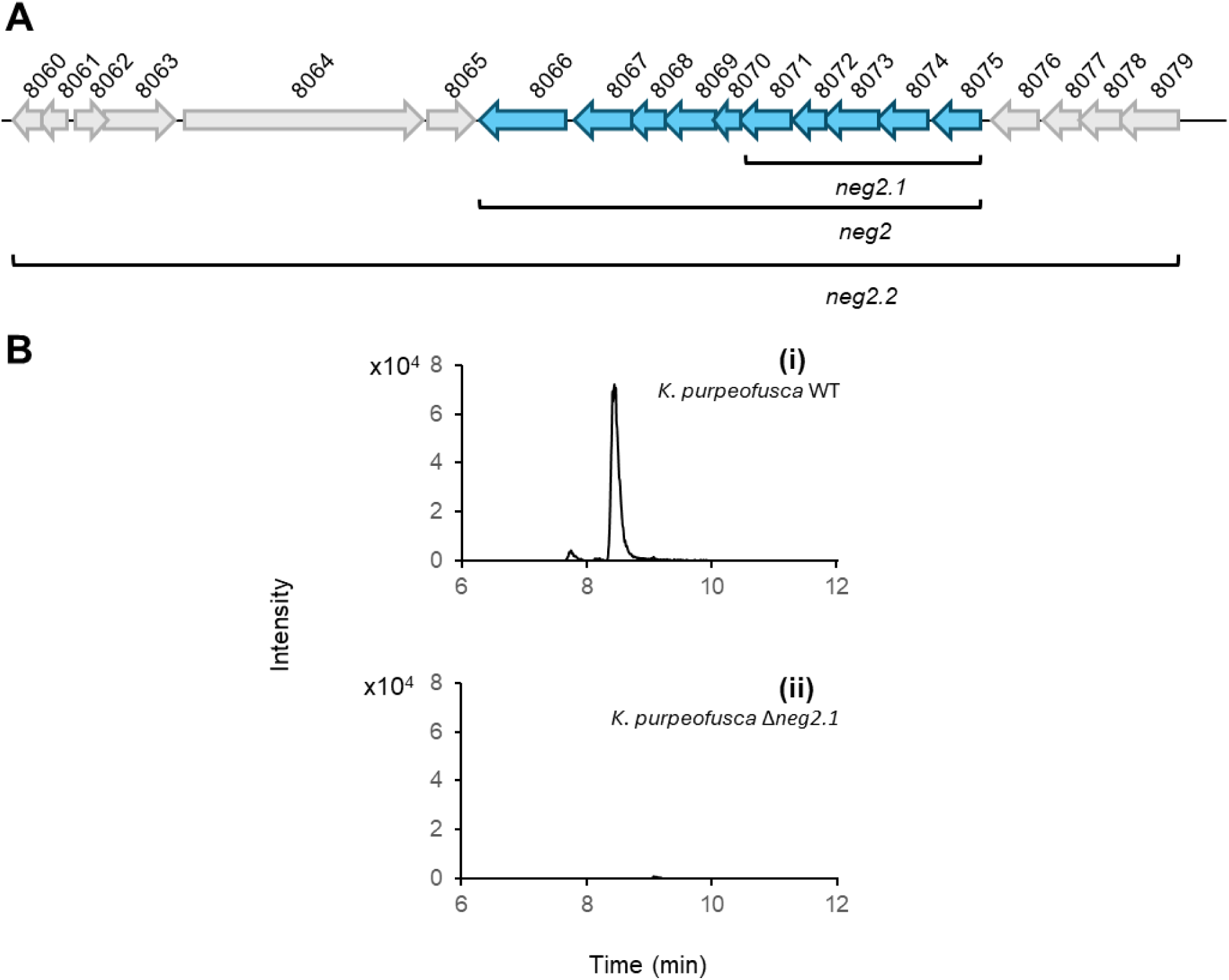
Genomic region putatively involved in negamycin biosynthesis (A) and LC-HRMS analysis of negamycin production (B). A: genes highlighted in blue correspond to the *neg2* region*. neg2.1* corresponds to the region deleted in the *K. purpeofusca* Δ*neg2.1* mutant. *neg2.2* corresponds to the insert of the BAC used for the heterologous expression in *S. albidoflavus* Del14. KamA2 (lysine aminomutase) is encoded by KPATCC21470_8074. B: Extracted ion chromatogram (m/z 471.2238 [M+H]^+^) of treated culture supernatants (see STAR Methods) of (i) *K. purpeofusca* ATCC 21470 (ii) *K. purpeofusca* Δ*neg2.1*. See also Table S2.

To test the involvement of the *neg2* region in negamycin biosynthesis, we generated a targeted deletion mutant of the KPATCC21470_8071–8075 genes (region *neg2.1*, Figure 4, A) using the classical two-step homologous recombination strategy. This deletion completely abolished negamycin production, implicating this region in the biosynthesis of the compound (Figure 4, B).

### Expression of *neg1* and *neg2.2* leads to negamycin production in S*treptomyces albidoflavus* Del14

In actinomycetes, BGCs responsible for natural product formation are typically co-localized within a single genomic region. The discovery that genes essential for negamycin biosynthesis are distributed across two separate loci is therefore unusual and prompted further investigation to functionally validate this split-cluster architecture.

To assess whether two regions are jointly sufficient for negamycin biosynthesis, we performed heterologous expression in engineered *Streptomyces* strains. The *creE-like* and *creD-like* genes from the *neg1* region were cloned into the integrative vector pIJ10257^25^ under the control of the constitutive *ermEp\** promoter. This construct (pIJ_*neg1*) integrates into the host genome via the ΦBT1 attachment site. In parallel, a BAC carrying *neg2.2* (KPATCC21470_8060-8079, ∼27 kb; Figure 4) was generated using the CAPTURE method (see STAR Methods). The resulting construct, BACn*eg2.2*, integrates site-specifically via the ΦC31 attachment site. By employing distinct integration systems, stable co-expression of both constructs can be achieved in a single host strain.

The two constructs were introduced individually and in combination into various heterologous *Streptomyces* hosts, including *Streptomyces lividans* TK24^26^, *Streptomyces coelicolor* M1146^27^, and *Streptomyces albidoflavus* Del14^28^ a mutant derived from *S. albus* J1074, recently reclassified as *S. albidoflavus*^29,30^. Remarkably, only the co-expression of pIJ_*neg1* and BAC*neg2.2* in *S. albidoflavus* Del14 resulted in production of negamycin, at levels comparable to the ones achieved with *K. purpeofusca*, as confirmed by LC-HRMS analysis (Figure 5). Neither construct alone, nor any other host-background combination, led to detectable levels of the compound.

**Figure 5:**
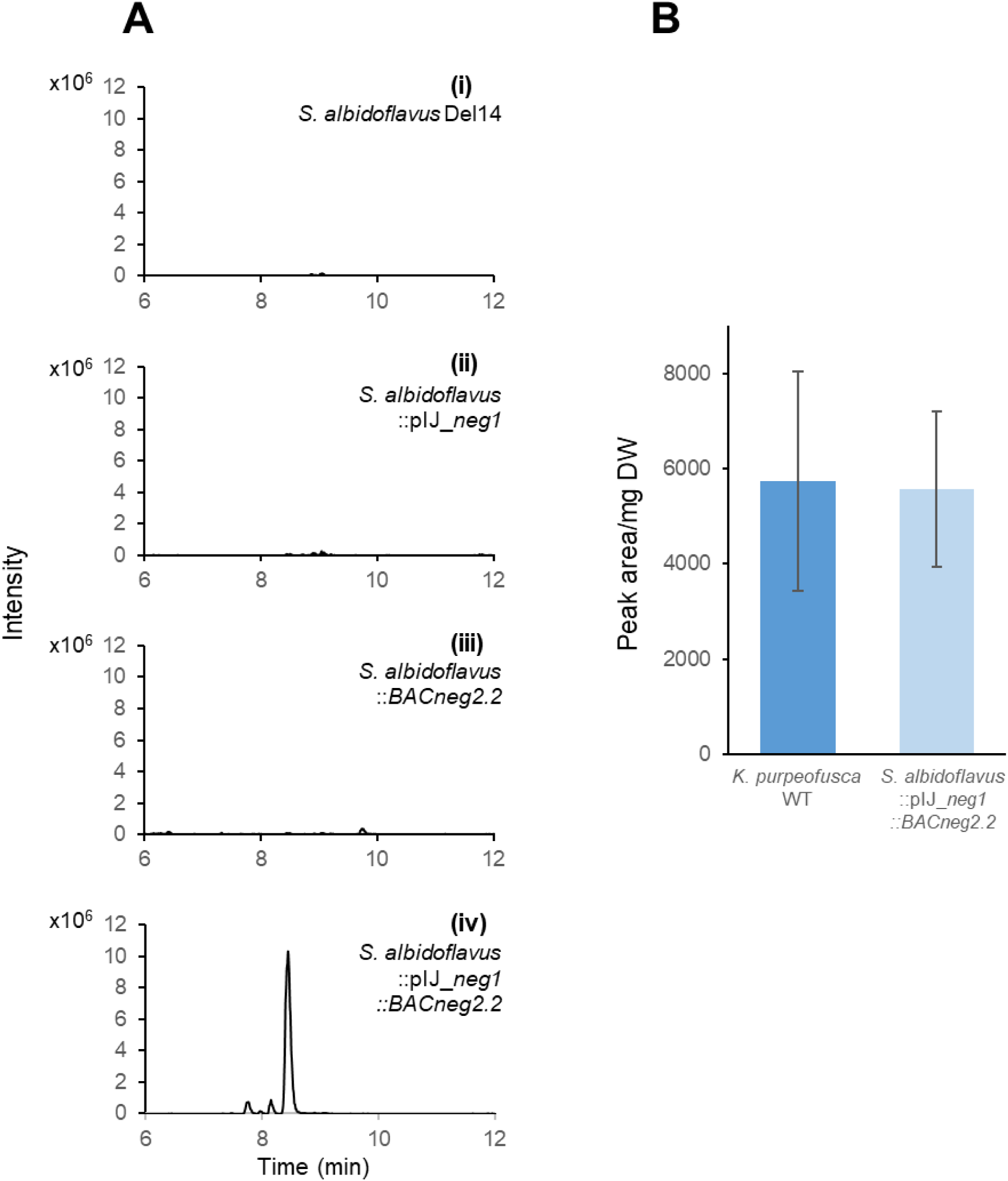
Negamycin production in the heterologous host *S. albidoflavus* Del14. A. Extracted ion chromatogram (*m/z* 471.2238 [M+H]^+^) of treated culture supernatants (see STAR Methods) of (i) *S. albidoflavus* Del14; (ii) *S. albidoflavus*::pIJ_*neg1*; (iii) *S. albidoflavus*::*BACneg2.2* and (iv) *S. albidoflavus*::pIJ_*neg1::BACneg2.2* B: Normalized negamycin production in *K. purpeofusca* WT and the heterologous host *S. albidoflavus* Del14 containing the *neg1* and *neg2.2.* Data are represented as mean ± SEM

These findings provide conclusive evidence that both *neg1 (creE*- and *creD*-like), encoding the nitrite-generating enzymes, and *neg2.2*, encompassing genes involved in β-lysine biosynthesis and likely N-N bond formation, are essential for negamycin production. Their combined expression led to the reconstitution of the complete biosynthetic pathway in a heterologous host, thus validating the functional interdependence of these two physically separated genomic loci.

Consistent with the findings of Wang et al.^20^, we identified two separate genomic loci *neg1*, and *neg2* as essential for negamycin biosynthesis. The *neg1* region contains *creE*- and *creD*-like genes likely involved in nitrite formation, while the *neg2* region (*negA–negJ*) encodes enzymes required for core scaffold assembly (Figure 6, Table 1). The pathway proposed by Wang et al.^20^ offers a valuable conceptual framework that complements our findings and provides a basis for future functional studies.

**Figure 6:**
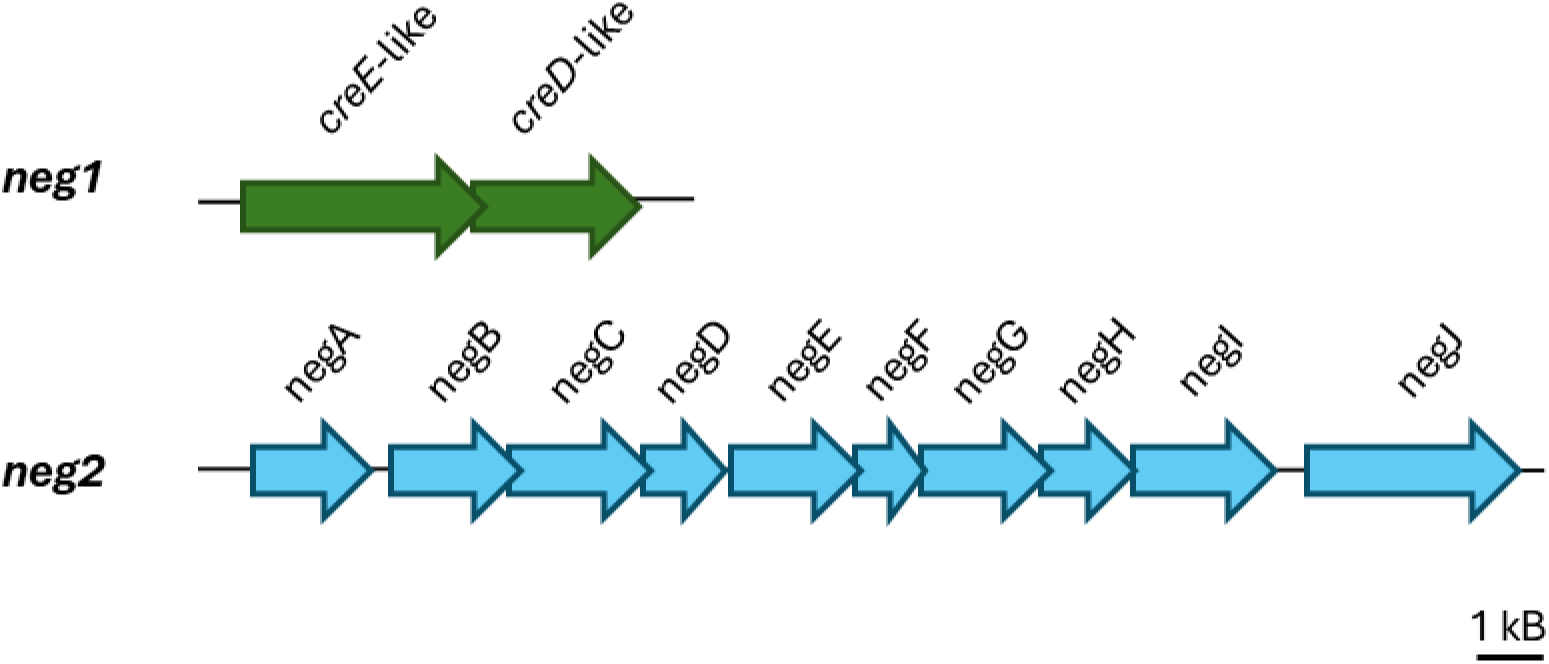
Split biosynthetic gene cluster involved in negamycin biosynthesis.

**Table 1:** Genes involved in negamycin biosynthesis and their putative function.

| ID<br>(KPATCC<br>21470) | name | putative function | Best hit in BLASTx results |  |  |
| --- | --- | --- | --- | --- | --- |
|  |  |  | query<br>cover<br>[%] | identity<br>[%] | protein ID |
| 4346 | *neg1_E* | flavin-dependent<br>monooxygenase | 93 | 99 | XOW80572.1 |
| **4347** | *neg1_D* | lyase family protein | 100 | 99 | WP_405005317 |
| **8075** | *negA* | serine hydrolase domain-containing protein | 100 | 98 | WP_340634362.1 |
| **8074** | *negB* | lysine-2,3-aminomutase-like protein | 91 | 99 | WP_266272324.1 |
| **8073** | *negC* | hypothetical protein | 100 | 99 | WP_266272325.1 |
| **8072** | *negD* | class I SAM-dependent methyltransferase | 100 | 99 | WP_266272326.1 |
| **8071** | *negE* | peptide ligase PGM-1 related protein | 100 | 99 | WP_266272327.1 |
| **8070** | *negF* | SET domain-containing protein-lysine N-methyltransferase | 99 | 98 | WP_266314959.1 |
| **8069** | *negG* | GNAT family N-acetyltransferase | 100 | 93 | WP_327075131.1 |
| **8068** | *negH* | alpha/beta hydrolase | 100 | 99 | WP_266272330.1 |
| **8067** | *negI* | MFS- transporter | 89 | 99 | WP_266272331.1 |
| **8066** | *negJ* | hypothetical protein | 100 | 99 | WP_266272332.1 |

## DISCUSSION

Our study began with an unexpected finding that deletion of the *neg1.1* (KPATCC21470_4344-4347), encoding the nitrite-generating homologs CreE and CreD, did not abolish negamycin production but reduced it only to approximately 9% of wild-type levels. This residual biosynthesis was traced back to low-level nitrite contamination in complex media components, such as soy flour and yeast extract, underscoring the critical role of nitrite as an essential biosynthetic precursor and highlighting how even minimal exogenous sources can mask the phenotypic effects of gene deletions. This observation, together with the feeding experiments using ^15^N labeled nitrite, underscores the critical importance of nitrite as a biosynthetic precursor and reveals how minimal exogenous contributions can obscure the effects of pathway deletions.

Another surprising finding was that not all genes within the *neg1.1* region were transcribed, notably the *creM*-like gene encoding a ligase. This was particularly unexpected given that the genes KPATCC21470_4344-4347 overlap. Correspondingly, CreM was absent in proteomic analyses. These observations suggested that additional genetic elements outside *neg1.1* contribute to negamycin biosynthesis. However, the lack of transcription of *creM*, despite genomic overlap with transcribed genes such as *creD*, points to the existence of complex regulatory mechanisms that are not yet understood and represent a limitation of our current study. Indeed, our integrated proteomic and transcriptomic analyses identified a second, physically separated chromosomal locus, *neg2*, which encodes enzymes responsible for β-lysine biosynthesis and likely subsequent scaffold assembly and N-N bond formation.

While split BGCs remain uncommon, several compelling examples suggest that such genomic organization may reflect modular evolution, regulatory decoupling, or cofactor recruitment strategies^31^. For example, in *Streptomyces* sp. MBT76, the biosynthesis of the catecholate-hydroxamate siderophore qinichelin is encoded by multiple distinct genomic loci. The main qinichelin BGC encodes core assembly functions, while physically distant BGCs provide essential precursors such as 3,4-dihydroxybenzoate and ornithine. This modular genomic organization highlights the complexity of precursor supply and suggests evolutionary advantages in distributing biosynthetic functions across the genome^32^. In *Streptomyces virginiae*, biosynthesis of griseoviridin and virginiamycin M is coordinated across separated BGCs that encode functionally interacting pathways^33^, as also observed for the biosyntheses of distamycin, disgocidine, and congocidine in *Streptomyces netropsis*^34^. In contrast to these examples, the two loci involved in negamycin biosynthesis do not encode any identifiable transcriptional regulators, raising the question of how gene expression is coordinated between the two regions. Future studies will be needed to investigate whether nitrite itself might play a regulatory role in coordinating this expression.

The non-colocalized genomic organization of *neg1* and *neg2* prompted rigorous validation through deletion, complementation, transcriptomic, and proteomic analyses, all confirming the essentiality of both loci. Furthermore, heterologous co-expression of *neg1* and *neg2* in *Streptomyces albidoflavus* Del14 successfully reconstituted negamycin biosynthesis, establishing their sufficiency.

During the final stages of our study, Wang et al.^20^ independently reported the identification of the exact same two biosynthetic loci in *K. purpeofusca*. Their work included the biochemical characterization of the heme-dependent monooxygenase NegJ, which catalyzes the formation of a hydrazine intermediate using nitrite and lysine-derived precursors.

Crucially, they also identified essential accessory proteins, a [2Fe-2S] ferredoxin (SpFd6) and a yet unidentified flavin reductase, required for NegJ activity, located somewhere else in the genome. Our transcriptomics data confirmed the expression of the KPUR_ATCC21470_5276 gene encoding the SpFd6 ferredoxin (Figure S5). These findings explain our own observation that heterologous production was successful only in *S. albidoflavus* Del14, a host that encodes compatible electron-transfer partners, which are missing in the other heterologous hosts. These findings highlight an additional layer of biosynthetic decentralization, consistent with observations in other natural product pathways such as pyochelin or aerobactin, where critical accessory proteins involved in iron acquisition or redox balancing are located outside core BGCs^35^. This underscores the necessity of considering broader genomic context and host redox and cofactor compatibility, when reconstituting natural product pathways in heterologous systems. Our findings suggest that distributed cofactor dependencies may be more common than previously appreciated and should be a key focus in future synthetic biology and metabolic engineering efforts.

Significantly, the detailed biosynthetic pathway for negamycin proposed by Wang et al.^20^, outlining sequential enzymatic steps culminating in hydrazide formation, is independently supported by our genetic, transcriptomic, and biochemical data. The convergent evidence concerning the roles of *creE/creD*-like in nitrite supply, β-lysine biosynthesis genes, and cofactor-supplying proteins substantially strengthens the validity of this comprehensive biosynthetic model.

Importantly, the heterologous expression of these genes led to negamycin production at levels comparable to the wild-type strain. This represents a key milestone and provides a solid foundation for further pathway optimization, both in terms of yield improvement and the generation of novel negamycin derivatives.

## RESOURCE AVAILABILITY

### Lead contact

Further information and requests for resources and reagents should be directed to and will be fulfilled by the lead contact, Sylvie Lautru.

### Materials availability

This study did not generate new unique reagents.

Plasmid and strains generated in this study will be shared upon request to Sylvie Lautru and Evi Stegmann and may require a completed material transfer agreement.

### Data and code availability

- The genome sequence of *K. purpeofusca* ATCC 21470 has been deposited at the EMBL database under the accession number GCA_965443725 (study number PRJEB89693 and sample number ERS24604253) and is publicly available as of the date of publication.
- The PacBio reads have been deposited at the EMBL database under the accession numbers ERR15007575; ERR15007574; ERR15007573 and are publicly available as of the date of publication.
- All RNA-seq Illumina read files as well as the raw counts have been deposited in NCBI’s Gene Expression Omnibus and are accessible under accession number
- The mass spectrometry proteomics data have been deposited to the ProteomeXchange Consortium via the PRIDE47 partner repository with the dataset identifier PXD064051.
- Any additional information required to reanalyze the data reported in this paper is available from the corresponding authors upon request.

## ACKNOWLEDGMENTS

This project is funded by the ANR (ANR-23-CE44-0042-01) and DFG STE 2052/4-1.NGS sequencing was performed with the support of the Core Facility for Genomics (“NCCT”) and the Institute for Medical Microbiology and Hygiene, University Hospital Tübingen. Data management and storage of raw data for this project were supported by the Quantitative Biology Center (QBiC), University of Tübingen, Germany. C.H. and H.B.O. acknowledge infrastructural support from the Cluster of Excellence EXC 2124: Controlling Microbes to Fight Infection (CMFI, project ID 390838134). E.S., HBO. and A:B. are grateful for funding by the German Center for Infection Research (DZIF). We thank Alessa Sauter for her work in this project during her master thesis. We thank Irina Droste-Borel for her technical support in LC-MSMS sample preparation. We thank Jean-Luc Pernodet for critical reading of the manuscript.

## AUTHOR CONTRIBUTIONS

Conceptualization, E.S. and S.L.; methodology and investigation, E.M. and H.S.O. identified the candidate loci. A.S generated most of the deletion mutants, complemented strains and heterologous hosts used in this study, performed transcriptomics (together with T.H. and K.N.), proteomics (together with M.F.-W. and B.M), RT-PCR experiments and Griess assay. C.H. established Fmoc derivatization protocol for LC-MS analysis of negamycin, C.G. established the Greiss assay protocol. S.T. did feeding experiments, final cultivations and LC-HR-MS^2^ analyses together with S.B. and D.T.; writing of the original draft, A.S. and S.T. wrote their respective parts; writing, review and editing, E.S. S.L. A.S., E.D. S.T.; funding acquisition, E.S. and S.L.; resources *K. purpeofusca* ATCC21470 WT and its genome sequence was provided by H.B.-O. and A.B.; supervision, E.S., S.L. and E.D.

## DECLARATION OF GENERATIVE AI AND AI-ASSISTED TECHNOLOGIES

During the preparation of this work, the author(s) used ChatGPT (GPT-4, accessed May 2025) in order to improve scientific writing of the manuscript. After using this tool or service, the authors reviewed and edited the content as needed and take full responsibility for the content of the publication.

## STAR★METHODS

KEY RESOURCES TABLE

KEY RESOURCES TABLE
| REAGENT OR RESOURCE | SOURCE | IDENTIFIER |
| --- | --- | --- |
| **Bacterial and virus strains** |  |  |
| See table S4 | This work |  |
| **Chemicals, peptides, and recombinant proteins** |  |  |
| Fmoc | Sigma-Aldrich | CAS#28920-43-6 |
| $^{15}\text{NO}_2^-$ | Cortecnec | CAS#1432-45-8 |
| L-Lysine-$^{13}\text{C}_6$ | Sigma-Aldrich | CAS#1228077-86-8 |
| Glycine-$^{15}\text{N}$ | Sigma-Aldrich | CAS#7299-33-4 |
| Q5® Hot Start High-Fidelity DNA Polymerase | NEB | M0491L |
| Taq DNA Polymerase | Thermo-Fisher | EP0401 |
| *NdeI* | Thermo Fisher | ER0585 |
| *HindIII* | Thermo Fisher | ER0501 |
| *EcoRI* | Thermo Fisher | ER0271 |
| *Apramycin* | Roth | M3450.0005 |
| Fosfomycin | Roth | 26016-99-9 |
| Nalidixid Acid | Roth | 389-08-2 |
| Hygromycin | Roth | 31282-04-9 |
| Chloramphenicol | Roth | 56-75-7 |
| TSB | Sigma Aldrich | 1.40143.5000 |
| **Deposited data** |  |  |
| Genome sequence and annotation | This work | GCA_965443725 |
| PacBio Reads |  | ERR15007573<br>ERR15007574<br>ERR15007575 |
| RNA-Seq Illumina read files as well as the raw counts | This work |  |
| Mass spectrometry proteomics data | This work | PXD064051 |
| **Oligonucleotides** |  |  |
| See table S5 for cloning primers | This work |  |
| **Recombinant DNA** |  |  |
| See Table S6 for plasmids | This Work |  |
| **Software and algorithms** |  |  |
| AntiSMASH 7.0 | https://antismash.secondarymetabolites.org/ |  |
| RAST | https://rast.nmpdr.org/rast.cgi |  |
| blastP | https://blast.ncbi.nlm.nih.gov/Blast.cgi |  |
| CluSeek 1.4.26 | https://lab111.mbu.cas.cz/cluseek/ |  |

| Other |  |  |
| --- | --- | --- |
| MCX Cartridges Waters OASIS ® | WATERS™ | SKU: 186000252 |

## EXPERIMENTAL MODEL AND STUDY PARTICIPANT DETAILS

All bacterial strains and plasmids used in this study are described in Tables S4 and S6.

## METHOD DETAILS

### Chemicals and reagents

Unless otherwise indicated, chemicals and enzymes were purchased from Sigma-Aldrich, Thermo-Fisher, and New England Biolabs. Oligonucleotides were synthesized by Eurofins Scientific Sigma Aldrich or IDT.

### Bacterial strains, plasmids and growth conditions

All bacterial strains and plasmids used in this study are listed in Tables S4 and S6, respectively. *E. coli* strains were cultivated in liquid or solid LB Medium at 37 °C^36^ using a selection marker (apramycin (100 µg/mL), kanamycin (100 µg/mL), chloramphenicol (50 µg/mL), hygromycin (50 µg/mL)) when necessary. *K. purpeofusca* strains were cultivated at 29 °C either in TSB medium using selection marker (fosfomycin (100 µg/mL), apramycin (100 µg/mL), hygromycin (50 µg/mL)) when necessary for 2 days and 180 rpm or on MS agar^37^ for genetic manipulation and spore stock preparation or Oatmeal agar (3,6% oats, 1.6% agar (w/v)).

For negamycin production, *K. purpeofusca* strains were cultivated in negamycin production medium (NPM)^1^ All strains were inoculated from spore stocks in 100 mL NPM and incubated for 6 days at 29 °C and 180 rpm.

### Preparation and manipulation of DNA

All oligonucleotides used in this study are listed in Table S5. *E. coli* transformations, *E. coli*/*K. purpeofusca* and *E. col*i/*Streptomyces* conjugations were performed under standard conditions^37^, either using mycelial conjugation or spore conjugation. DNA sequencing was performed by Sanger sequencing at Eurofins.

### *K. purpeofusca* genome sequencing, assembly and annotation

High molecular weight genomic DNA was extracted using the NucleoBond® HMW DNA kit (Macherey-Nagel, Düren, Germany; Cat.-no-: 740160.20) according to the manufacturer’s protocol. The genome was sequenced by the Magrogen company using a PacBio2 RSII instrument, generating 137 431 reads that were assembled using HGAP3, yielding two contigs. The gap between these contigs was filled by amplifying and sequencing a PCR product using oligonucleotides 1 and 2. The genome sequence was annotated using the RAST annotation server^38^. The ENA accession number of the genome is GCA_965443725.

### Generating competent cells

DNA was transferred either into *E. coli* via transformation or in Actinomycetes via conjugation. For transformation, either self-prepared chemo competent *E. coli*^39^ or electrocompetent cells^40^ were used.

### Molecular cloning

In this study, different methods for DNA modification were used. For amplification of DNA fragments Polymerase chain reactions (PCR) were performed using “Q5 Hot Start High-Fidelity DNA Polymerase” or “Taq DNA Polymerase”. For linearization of pGusA21, pRM4 and pIJ12570 vector DNA the restriction endonucleases *Nde*I and *Hin*dIII were used.

#### Construction of plasmids used in this study

The plasmids used in this work for gene deletion, complementation, and heterologous expression, were constructed primarily using standard Gibson assembly method. For plasmid assembly, all constructs were transferred to *E. coli* NovaBlue.

**pGusA21_Δ*neg1.1*** – Used for the deletion of the *neg1.1* region (KP_ATCC21470_4344 - _4347) in *K. purpeofusca*. The plasmid was constructed via restriction cloning by amplifying 1500 bp flanking regions using the oligonucleotide pairs 3/4 & 5/6 (Table S5). The fragments were cloned into the vector pGusA21 using *Nde*I, *Hin*dIII, and *Eco*RI restriction sites.

**pGusA21_Δ*neg2.1*** – Used for the deletion of *neg2.1* (KP_ATCC21470_8071 – 8075) in *K. purpeofusca*. The 1500 bp flanking regions were amplified with specific oligonucleotide pairs 15/16 & 17/18 (Table S5) and assembled into the with *Nde*I and *Hin*dIII linearized vector pGusA21.

**pRM4_*neg1.2*** – Used for the complementation of *neg1.1* in *K. purpeofusca_Δneg1.1*. The insert was amplified in three fragments using oligonucleotide pairs 9/10, 11/12 and 13/14 (Table S5) to ensure correct gene orientation while excluding KPATCC21470_4341 postulated to encode a negative regulator. **pIJ_*neg1*** – Used for the heterologous expression of *creE*-like and *creD-*like. The genes *creE* and *creD* (KPATCC21470_4346 and 4347) were amplified in one fragment using oligonucleotide pair 35/36 and cloned int the vector pIJ10257 (Table S5).

#### Cloning of BACneg2.2

To heterologously express the *neg2.2* region (KPATCC21470_8066–8075), the BAC*neg2.2* construct was cloned using the CAPTURE method—a Cas12a-assisted, precise targeted cloning technique that employs *in vivo* Cre-lox recombination^40^.

For the preparation of high molecular weight genomic DNA (gDNA), *Kitasatospora purpeofusca* was cultured in 50 mL ISP2 medium at 29 °C with shaking at 180 rpm for 2 days. The extracted DNA was stored at 4 °C without freezing.

To isolate the 27 kb *neg2.2* region (KPATCC21470_8060–_8079) from the genome, sgRNAs 27 and 28 (Table S5) were utilized. For BAC*neg2.2* cloning, the upstream receiver fragment was amplified using primer pair 23/26 with pBE48 as the template, while the downstream receiver was amplified using primer pair 24/25 with pBE45 as the template.

### Generation of *K. purpeofusca* deletion mutants

To generate in-frame deletion mutants of *Kitasatospora purpeofusca*, the non-replicative vector pGusA21 was used. Flanking homologous regions (∼1.5 kb) of the target loci *neg1.1* and *neg2.1* were cloned into pGusA21, resulting in the construction of the deletion plasmids pGusA21_Δneg1.1 and pGusA21_Δneg2.1.

Each plasmid was introduced into *K. purpeofusca* via mycelial conjugation, yielding recombinant strains. Successful single-crossover integration of the deletion plasmids into the genome was confirmed by blue-white screening and selection on apramycin-containing media.

Recombinant strains harboring either pGusA21_Δ*neg1.1* or pGusA21_Δ*neg2.1* were subjected to a stress-based protocol using S-medium (composition: 4 g peptone, 4 g yeast extract, 4 g K₂HPO₄, 2 g KH₂PO₄, 10 g glycine in 800 mL ddH₂O; after autoclaving, supplemented with 10 g glucose and 0.5 g MgSO₄·7H₂O in 200 mL ddH₂O) containing apramycin. This approach was aimed at increasing the frequency of double-crossover events. Subsequently, protoplasts were generated to ensure isolation of single clones.

The resulting protoplasts were plated on R5 agar containing x-Gluc for regeneration and blue-white selection, allowing identification of colonies that had lost the deletion plasmid. White colonies were then screened by PCR using oligonucleotide pairs 7/8 or 19/20 (Table S5) to confirm successful in-frame deletion of the respective target regions.

### Complementation of deletion mutant *K. purpeofusca* Δ*neg1.1*

For the genetic complementation of the deletion mutant *K. purpeofusca* Δ*neg1.1,* the non-replicative plasmid pRM4_*neg1.2*, containing region *neg1.2* was used for complementing *K. purpeofusca* Δ*neg1.1.* The deletion mutant *K. purpeofusca* Δ*neg1.1.*was transformed with the appropriate plasmid via mycelial conjugation. Successful integration of pRM4_*neg1.2* into genome was verified both via growing with selection marker apramycin and via PCR using oligonucleotide pair 21/22 (Table S5).

### Generation of heterologous hosts

To heterologously express the negamycin biosynthetic genes, the plasmid pIJ_*neg1* was constructed via Gibson assembly, and the BAC*neg2.2* construct was generated using the CAPTURE method.

*Streptomyces lividans* TK24, *S. coelicolor* M1154, and *S. albus* Del14 were initially transformed with pIJ_*neg1* through spore conjugation. Successful integration of the plasmid was confirmed by growth on hygromycin-containing media and PCR verification using specific oligonucleotide pair 44/45 (Table S5).

The verified heterologous host strains carrying pIJ_*neg1* were subsequently transformed with BAC*neg2.2* via spore conjugation. Integration was verified by selection on media containing both hygromycin and apramycin, as well as by PCR using oligonucleotide pairs 29/30, 31/32, and 33/34.

In parallel, the same strains (*S. lividans* TK24, *S. coelicolor* M1154, and *S. albidoflavus* Del14) were independently transformed with BAC_neg2.2 following the same procedure.

### Culture supernatant purification and Fmoc derivatization

*K. purpeofusca* ATCC21470 and its derivatives were cultivated in NPM medium at 30 °C for 8 days. 10 mL of culture supernatant were partially purified using Oasis MCX cartridges (strong cation exchanger, 60 µm, 60 mg/3 mL; Waters). Cartridges were conditioned with 5 mL of methanol, followed by equilibration with 5 mL of 2% formic acid. The culture supernatant was diluted in 40 mL of 2% formic acid before loading onto the cartridge. After loading, the cartridges were washed sequentially with 5 mL of 2% formic acid, followed by two washes with 5 mL of methanol to remove hydrophobic compounds. Negamycin was finally eluted in two 5 mL fractions of 5% ammonium hydroxide (NH₄OH). The eluates were lyophilized and resuspended in 2 mL of water.

For detection by high-resolution LC-MS (LC-HRMS), samples were derivatized with Fluorenylmethyloxycarbonyl group (Fmoc). The reaction was carried out by mixing 200 µL of sample, 200 µL of borate buffer (crystallized boric acid, R.P. Normapur, 200 mM, pH 8.8), and 100 µL of Fmoc chloride (Sigma-Aldrich, 10 mM in acetonitrile). The mixture was incubated at room temperature for 30 min, diluted 1:1 in 0.1% formic acid, and analyzed by LC-HRMS.

### LC-HRMS/MS analysis for negamycin production

Liquid chromatography-tandem mass spectrometry with high resolution (LC-HRMS/MS) experiments were performed on 1260, Infinity II LC system (Agilent Technologies, France) coupled to a timsTOF mass spectrometer (Bruker, France) equipped with an electrospray ionization (ESI) source operated in the positive mode. The chromatographic separation was carried out on a Poroshell 120 SB C18 (2.1×150 mm 2.7 μm) column heated at 40 °C, the flow rate was set at 400 μL/min with water as solvent A and acetonitrile as solvent B with 0.1% of formic acid in A and B. 10 µL of sample were injected using the following gradient: 0 min (5% B), 10 min (50% B), held for 10 minutes, at 20 min (100% B) held for 5 min and 9 minutes of equilibration time (5% B).

Ion source parameters were set as follows: capillary voltage 4.2 kV, nebulizer 3.0 bar, dry gas 10.0 L/min, desolvatation temperature 250°C no collision in source. Nitrogen was used as drying gas in the source and for collision experiments. The instrument was daily calibrated in the *m/z* 20-1350 range using ESI-L low Concentration Tuning Mix solution (Agilent Technology). For Auto MS/MS experiments, parameters were set as follows: precursor ion range *m/z* 99-1350, No. of precursor 5, spectral rate 10.0 Hz, CID between 10 eV to 50 eV.

The elemental compositions for all ions were determined with the instrument software DataAnalysis, the precision mass measurement was less than 5 ppm.

### Feeding experiments with ^15^N-labeled nitrite

100mL cultures of *K. purpeofusca* were inoculated in NPM with 10^6^ spores of either the *K. purpeofusca* wild type (WT) strain or the Δ*neg1* mutant and incubated at 30°C for 8 days. ^15^N**-**labeled sodium nitrite (^15^NO2^−^, purchased from CortecNet) was added to the cultures at a final concentration of 0.5 mM at t = 0, 12, 24, 48, and 96 h. After incubation, the supernatants were processed as previously described prior to LC-HR-MS analysis.

### Feeding experiments with labeled lysine and glycine

Cultures of *K. purpeofusca* wild-type were grown in 100 mL of NPM medium inoculated with 10⁶ spores and incubated at 28 °C for 10 days. ¹³C₆ -labeled L-lysine, ¹³C₂-labeled glycine, and ¹⁵N-labeled glycine (all purchased from Sigma-Aldrich) were each added at a final concentration of 1 mM at t = 0, 48, 60, 72, and 84 h. After incubation, culture supernatants were processed as previously described prior to LC-HRMS analysis.

### Expression of negamycin biosynthetic genes

#### Transcriptomics

To analyze the expression levels in *K. purpeofusca* WT, transcriptomics was performed.

The RNA was isolated after 4 days of cultivation under negamycin production conditions using TRizol^41^ analogous to the protocol provided by Thermo scientific.

#### RNA library preparation and sequencing

The total RNA was quantified with a Qubit RNA BR Assay Kit (Thermo Fisher) and RNA integrity was checked on Agilent 2100 BioAnalyzer with RNA 6000 Pico Kit (Agilent).

Library preparation for RNA-seq was performed with Illumina Stranded Total RNA Prep, Ligation with Ribo-Zero Plus Microbiome according to the manufacturer’s instructions. In brief, 100 ng of total RNA per sample were subjected to rRNA depletion, followed by cDNA library construction, adapter ligation and 15 cycles of barcoding PCR. Obtained libraries were quantified with Qubit 1x DNA HS Assay Kit (Thermo Fisher) and the fragment distribution was checked Agilent 2100 BioAnalyzer using High Sensitivity DNA Kit (Agilent). Libraries were subsequently pooled and sequenced on an Illumina NextSeq 550 device using NextSeq 500 High Output Kit v2.5 (75 cycles) (Illumina) with a run mode 72,10,10,0.

#### Demultiplexing and quality control

The sequencing was demultiplexed with the latest version of the nextflow pipeline: <u>nf-core/demultiplex</u>. For demultiplexing bcl2fastq was used and the quality was checked with fastp.

Sequencing statistics, including the quality per base and adapter content assessment, were conducted with FastQC (v0.11.8).^42^ All reads mappings were performed against the genome of GCA_965443725. The mappings of all samples were conducted with HISAT2 (v2.1.0)^43^, using the following parameters: spliced alignment of reads was disabled, and strand-specific information was set to reverse complemented (HISAT2 parameter --no-spliced-alignment and --rna-strandness “R”). The resulting mapping files in SAM format were converted to BAM format using SAMtools (v1.9)^44^. Mapping statistics, including strand specificity estimation and percentage of mapped reads, were conducted with the RNA-Seq module of QualiMap2 (v2.2.2-a)^45^. Gene counts for all samples were computed with featureCounts (v1.6.4)^46^, where the selected feature type was set to transcript records (featureCounts parameter -t transcript). A quality check for ribosomal rRNA was performed with a self-written script based on the absolute counts of annotated rRNAs. To assess variability across the replicates of each time series, a principal component analysis (PCA) was conducted with the DESeq2 package (v1.28.1)^47^. For the computation of genes differentially expressed between the mutants, DESeq2 was applied to the absolute gene counts as computed with featureCounts. For differences between the Δ*neg1* mutant and the wildtype strain, genes with an adjusted p-value (FDR) < 0.05 and absolute log2 fold change (FC) > 1 were reported as differentially expressed.

#### Transcriptional analysis

To analyze the expression of negamycin biosynthetic genes in *K. purpeofusca* WT after different time points of cultivation transcriptional analysis was performed. *K. purpeofusca* WT was cultivated under negamycin production conditions. RNA isolation was performed after 24 h, 36 h, and 48 h using the Kit RNeasy from Qiagen. DNase treatment was performed using DNase from Macherey Nagel. RNA was transcribed to cDNA, which was used for specific investigation of expression of specific negamycin biosynthetic genes using primers 27 - 55 (S5) for the amplification of fragments of the negamycin biosynthetic genes.

### Proteomics

In order to analyze the protein profile of *K. purpeofusca* WT and *K. purpeofusca* Δ*neg1.1* proteomics was performed. Both strains were cultivated under negamycin production conditions and proteins were purified after 72 h using SDS buffer.

#### Protein purification

The cell pellet was dissolved in SDS buffer (4% w/v sodium dodecyl sulfate (SDS), in 100 mM tris(hydroxymethyl)aminomethane (Tris)/HCl; pH 8.0), using 1 mL of buffer per 20–50 mg of cells and incubated at 95 °C for 10 min, with careful vortexing at intervals. Afterwards the suspension was chilled on ice for 2–5 min to prevent protein degradation. To shear the DNA and reduce viscosity, the suspension was sonicated for 30–300 sec until it reached a watery consistency. Sonication was performed using a Branson Sonifier 250 with a 5 mm microtip, set to an output control of 5 and a duty cycle of 40%.

Next, Dithiothreitol ((DTT) 1 M) to a final concentration of 10 mM was added and incubated for 45 min at room temperature (RT) while shaking at 650 rpm. Iodoacetamide ((IAA) 0.55 M) to a final concentration of 5.5 mM was added and incubated for another 45 min at RT, while shaking at 650 rpm in the dark.

The suspension was then centrifuged for 15 min at 12,000 × g and the supernatant was carefully collected for protein precipitation. 1 volume of the supernatant was mixed with 8 volumes of ice-cold acetone and 1 volume of methanol. The mixture was vortexed thoroughly and incubated overnight at -20 °C.

The precipitated proteins were collected by centrifugation at 500–1000 × g for 5 min. The resulting pellet was washed by resuspending it in 80% (v/v) aqueous acetone at RT twice followed by air drying for 10-15 min at RT.

The dried protein pellet was rehydrated in denaturation buffer (6 M urea, 2 M thiourea in 10 mM Tris/HCl; pH 7.5), making it ready for subsequent analyses.

#### In-solution digestion, dimethylation labeling, LC-MSMS

20 µg of each sample were digested with trypsin using the Preomics iST Kit according to the instruction manual. After evaporation of the organic solvent, peptides were loaded on C18 stage tips^48^ and labeled with dimethyl “light” ((CH3)2) and dimethyl “medium-heavy” ((CH1D2)2), respectively as described elsewhere^49^. In brief: C18 bound peptides were flushed with 1 ml labeling solution and washed with 0.5 ml of HPLC solvent A (0.1% formic acid). Labeling efficiency tests were performed, and samples were pooled in equal ratio.

The peptide mixture was analysed on anEasy-nLC 1200 UHPLC system coupled to a quadrupole Orbitrap Exploris 480 mass spectrometer *via* a nanoelectrospray ion source (all Thermo Fisher Scientific) as described previously^50^ with slight modifications. Peptides were separated with an 87min segmented gradient of 10-33-50-90% of HPLC solvent B (80% acetonitrile in 0.1% formic acid) at a flow rate of 200 nl/min. MS and MS/MS spectra were recorded with a resolution of 60,000 and 30,000, respectively, whereby AGC was set to standard and fill time was set to automatic. The 20 most intense precursor ions were picked for HCD fragmentation in each scan cycle. Masses were excluded from further fragmentations for 30s.

#### MS data processing and statistical analysis

Acquired MS data were further analyzed using MaxQuant software version 2.5.0.0^51^ with integrated Andromeda search engine^52^.

Spectra were searched against a *K. purpeofusca* database from (1^st^ July 2025, 8,625 entries), and a database consisting of 285 common potential contaminants. Default search parameters were applied, except for multiplicity that was set to two with DimethylLys0/DimethylNter0 and DimethylLys4/DimethylNter4 for Light (L) and Heavy (H) labeling, respectively. The option “Re-quantify” was enabled.

Calculation of the significance B (psigB) for each protein ratio was performed with the Perseus software version 1.6.15.0^53^. Protein ratios were log2 transformed and blotted against log10 transformed sums of related protein intensities. Proteins with psigB < 0.05 were considered to be differentially expressed.

### Nitrite test by Griess reaction

To investigate whether the NPM contains nitrite, *G*riess-reaction^54^ was performed. The reagent was made by mixing 2.3 mL Phosphoric acid (85%), 1g Sulfanilamide, 0.1g Naphtylethylenediamine and 97.7ml water. Samples were prepared analogous to the protocol provided by Promega (Promega, Cat. No. G2930)

NPM itself and the medium components (soy flour and yeast extract) were tested. Instead of measuring the OD using a microplate reader, HPLC-MS was used to identify the ion of the resulting diazonium salt product (C18H20N5O2S^+^) m/z = 370.1332 [M+H]^+^

#### LC-HRMS analysis for Griess analysis

High resolution RP-LCMS runs were performed on an Agilent 1290 Infinity II system by Agilent Technologies (Waldbronn, Germany) coupled to a Bruker Impact II QTof mass spectrometer by Bruker Daltonics (Bremen, Germany). A gradient with (A) H2O + 0.1% formic acid and (B) acetonitrile + 0.1% formic acid was run on a Kinetex C18 1.7 µm core-shell 100 Å column (dimensions: 50 x 2.1 mm (l. X i.d.)) by Phenomenex (Torrance, CA, USA). The gradient was as follows: 0.0 min: 5% B, 10 min: 100% B, 12 min: 100% B, 13 min: 5% B, followed by 2 min reequilibration time.

