## Supplemental Tables and Figures for "Identification of the negamycin split biosynthetic gene cluster in *Kitasatospora purpeofusca* ATCC21470"

SUPPLEMENTAL INFORMATION

**Table S 1: Identification of candidate negamycin biosynthetic enzymes encoded in the genome of *K. purpeofusca*.**
Protein sequences of enzymes known to be involved in N-N bond and β-lysine biosynthesis were BLASTed against the CDS encoded within *K. purpeofusca* genome.

| | | | **Best hit in *K. purpeofusca* genome** | | |
|---|---|---|---|---|---|
| **protein number** | **accession name** | **function** | **ID (KPATCC21470_)** | **query cover [%]** | **identity [%]** |
| ALA99210.1 | CreM | long-chain fatty acid CoA ligase | 4344 | 97 | 49 |
| ALA99202.1 | CreE | oxidoreductase | 4346 | 93 | 60 |
| ALA99201.1 | CreD | 3-carboxy-cis,cis-muconate cycloisomerase | 4347 | 97 | 68 |
| BAW27702.1 | Spb38 | oxygenase | 8504 | 89 | 39 |
| BAW27703.1 | Spb39 | oxidoreductase | 8500 | 41 | 34 |
| BAW27704.1 | Spb40 | C-terminal MetRS-like domain | 8501 | 32 | 35 |
| WP_191740046.1 | KamA1 | lysine 2,3-aminomutase | 2073 | 74 | 31 |
| WP_191740046.1 | KamA2 | lysine 2,3-aminomutase | 8074 | 82 | 41 |
| WP_191740046.1 | KamA3 | lysine 2,3-aminomutase | 8180 | 92 | 48 |

**Table S 2: Predicted functions of genes within the *neg1.2* and *neg2.2* regions.**

| | | Best hit in BLASTx results | | |
|---|---|---|---|---|
| ***neg1.2* region** | | | | |
| **ID (KPATCC21470_)** | **putative function** | **query cover [%]** | **Identity [%]** | **protein ID** |
| **4339** | FAD-binding oxidoreductase | 100 | 99 | WP_266311934.1 |
| **4340** | VOC family protein | 99 | 99 | WP_380516065.1 |
| **4341** | TetR/AcrR family transcriptional regulator | 87 | 99 | WP_380440986.1 |
| **4342** | GNAT family N-acetyltransferase | 99 | 98 | WP_266270411.1 |
| **4343** | SRPBCC family protein | 99 | 99 | WP_266311938.1 |
| **4344** | class I adenylate-forming enzyme family protein | 100 | 99 | WP_435649838.1 |
| **4345** | FAD-dependent monooxygenase | 100 | 99 | WP_435649839.1 |
| **4346** | flavin-dependent monoxygenase | 93 | 99 | XOW80572.1 |
| **4347** | lyase family protein | 100 | 98 | WP_405005317 |
| ***neg2.2* region** | | | | |
| **8060** | phosphatase PAP2 family protein | 91 | 98 | WP_380517377.1 |
| **8061** | DedA family protein | 86 | 98 | WP_266272338.1 |
| **8062** | response regulator transcription factor | 100 | 99 | WP_380301093.1 |
| **8063** | sensor histidine kinase | 93 | 99 | WP_266272336.1 |
| **8064** | RHS repeat-associated core domain-containing protein | 99 | 99 | WP_435651888.1 |
| **8065** | hypothetical protein | 100 | 99 | WP_266272334.1 |
| **8066** | hypothetical protein | 100 | 99 | WP_266272332.1 |
| **8067** | MFS- transporter | 89 | 99 | WP_266272331.1 |
| **8068** | alpha/beta hydrolase | 100 | 98 | WP_266272330.1 |
| **8069** | GNAT family N-acetyltransferase | 100 | 93 | WP_327075131.1 |
| **8070** | SET domain-containing protein-lysine N-methyltransferase | 99 | 98 | WP_266314959.1 |
| **8071** | peptide ligase PGM-1 related protein | 100 | 99 | WP_266272327.1 |
| **8072** | class I SAM-dependent methyltransferase | 100 | 99 | WP_266272326.1 |
| **8073** | hypothetical protein | 100 | 99 | WP_266272325.1 |
| **8074** | lysine-2,3-aminomutase-like protein | 91 | 99 | WP_266272324.1 |
| **8075** | serine hydrolase domain-containing protein | 100 | 98 | WP_340634362.1 |
| **8076** | hydroxyacid dehydrogenase | 93 | 99 | WP_407661069.1 |

| | | | | |
|---|---|---|---|---|
| **8077** | carbohydrate ABC transporter permease | 100 | 99 | WP_380513544.1 |
| **8078** | carbohydrate ABC transporter permease | 73 | 99 | WP_266274541.1 |
| **8079** | ABC transporter substrate-binding protein | 90 | 99 | WP_380513546.1 |

**Table S 3: Proteomics; Proteins significantly more abundant in *K. purpeofusca* WT compared to *K. purpeofusca* ∆*neg1.1*.**

Proteins more abundant in *K. purpeofusca* WT compared to *K. purpeofusca* ∆*neg1.1* with significance of 0.01 and 0.05. proteins encoded by genes located in *neg2.1* are highlighted in bolt letters.

| protein ID (KPATCC 21470_) | hypothetical function | log2 fold change | significance B |
|---|---|---|---|
| 4451 | hypothetical protein | -2,7179 | 1,2E-07 |
| 5030 | peptidyl-prolyl cis-trans isomerase, cyclophilin type | -2,6564 | 2,4E-07 |
| 0687 | hypothetical protein | -2,0615 | 6,4E-05 |
| 7471 | hypothetical protein | -2,0310 | 6,6E-05 |
| 1888 | hypothetical protein | -2,0210 | 7,2E-05 |
| 1472 | putative neutral zinc metalloprotease | -1,9797 | 1,0E-04 |
| 8066 | hypothetical protein | -1,9495 | 1,3E-04 |
| 2708 | hypothetical protein | -1,9480 | 1,3E-04 |
| 6709 | rhodanese-related sulfurtransferase, 1 domain | -1,8825 | 2,6E-04 |
| 5276 | 4Fe-4S ferredoxin, iron-sulfur binding | -1,8824 | 2,7E-04 |
| 1153 | putative neutral zinc metalloprotease | -1,8801 | 2,7E-04 |
| 2297 | acetyltransferase, GNAT family | -1,8325 | 3,9E-04 |
| 8325 | formyltransferase | -1,8063 | 4,1E-04 |
| **8075** | **serine hydrolase domain-containing protein** | **-1,7955** | **4,4E-04** |
| 5517 | putative neutral zinc metalloprotease | -1,7505 | 7,0E-04 |
| 8542 | hypothetical protein | -1,7193 | 7,8E-04 |
| 1457 | hypothetical protein | -1,6939 | 9,4E-04 |
| **8073** | **hypothetical protein** | **-1,6911** | **1,1E-03** |
| 4280 | hypothetical protein | -1,6888 | 1,1E-03 |
| 2786 | putative hydroxylase | -1,6744 | 1,2E-03 |
| 7164 | PTS system, N-acetylglucosamine-specific IIA component | -1,6576 | 1,3E-03 |
| 6612 | isochorismat synthase siderophore | -1,6307 | 1,5E-03 |
| 1563 | N-acetylmuramoyl-L-alanine amidase | -1,6423 | 1,5E-03 |
| 7264 | sphingomyelinase | -1,6246 | 1,5E-03 |
| **8072** | **hypothetical protein** | **-1,6372** | **1,5E-03** |
| 1075 | hypothetical protein | -1,6364 | 1,6E-03 |
| 1122 | hypothetical protein | -1,6270 | 1,7E-03 |

| | | | |
|---|---|---|---|
| 6984 | hypothetical protein | -1,6123 | 1,7E-03 |
| **8074** | **NegB (KamA2)** | **-1,5806** | **2,3E-03** |
| 1402 | hypothetical protein | -1,5592 | 2,4E-03 |
| 2588 | putative large secreted protein | -1,5225 | 3,3E-03 |
| 2732 | hypothetical protein | -1,5025 | 3,5E-03 |
| 7508 | enoyl-CoA hydratase | -1,5006 | 3,5E-03 |
| 7406 | hypothetical protein | -1,4912 | 4,0E-03 |
| 1756 | hypothetical protein | -1,4592 | 4,9E-03 |
| 4448 | hypothetical protein | -1,4473 | 5,2E-03 |
| 8205 | hypothetical protein | -1,4353 | 5,4E-03 |
| 1373 | putative lyase PtlJ | -1,4142 | 6,1E-03 |
| 6256 | hypothetical protein | -1,3888 | 7,4E-03 |
| 3412 | organic hydroperoxide resistance protein | -1,3815 | 7,7E-03 |
| 7580 | hypothetical protein | -1,3648 | 8,2E-03 |
| 6632 | uncharacterized protein YehL | -1,3540 | 8,8E-03 |
| 6115 | hypothetical protein | -1,3388 | 9,6E-03 |
| 7822 | phospholipase C | -1,3344 | 1,0E-02 |
| 4220 | hypothetical protein | -1,3255 | 1,1E-02 |
| 7801 | polyphosphate kinase | -1,3201 | 1,1E-02 |
| 2754 | 2',3'-cyclic-nucleotide 2'-phosphodiesterase | -1,3110 | 1,2E-02 |
| 7911 | near of KamA2 upstream | -1,3088 | 1,2E-02 |
| 5523 | methylmalonyl-CoA mutase | -1,2973 | 1,2E-02 |
| 5021 | cold shock protein of CSP family | -1,2981 | 1,2E-02 |
| 4738 | putative hydroxylase | -1,2911 | 1,3E-02 |
| 3392 | phospholipase C | -1,2786 | 1,4E-02 |
| 4045 | putative secreted protease | -1,2765 | 1,4E-02 |
| 6012 | hypothetical protein | -1,2652 | 1,5E-02 |
| **8071** | **peptide ligase PGM1-related protein** | **-1,2634** | **1,5E-02** |
| 1573 | polyketide synthase modules and related proteins | -1,2358 | 1,7E-02 |
| 1130 | uncharacterized lipoprotein aminopeptidase LpqL | -1,2339 | 1,8E-02 |
| 2970 | glutamine synthetase adenylyl-L-tyrosine phosphorylase | -1,2331 | 1,8E-02 |

| | | | |
|---|---|---|---|
| 7929 | hypothetical protein | -1,2213 | 1,9E-02 |
| 4199 | 5'-nucleotidase | -1,1953 | 2,2E-02 |
| 7802 | phospholipase C | -1,1876 | 2,2E-02 |
| 4229 | RNA polymerase ECF-type sigma factor | -1,1310 | 3,0E-02 |
| 2966 | putative neutral zinc metalloprotease | -1,1251 | 3,1E-02 |
| 5172 | L,D-transpeptidase | -1,1250 | 3,1E-02 |
| 7266 | hypothetical protein | -1,0998 | 3,5E-02 |
| 0980 | putative secreted protein | -1,0950 | 3,6E-02 |
| 5519 | putative neutral zinc metalloprotease | -1,0720 | 4,0E-02 |
| 8187 | hypothetical protein | -1,0617 | 4,2E-02 |
| 0613 | peptidoglycan N-acetylglucosamine deacetylase | -1,0616 | 4,2E-02 |
| 1414 | hypothetical protein | -1,0613 | 4,2E-02 |
| 5940 | copper(I) chaperone CopZ | -1,0532 | 4,4E-02 |
| 2496 | long-chain-fatty-acid--CoA ligase | -1,0433 | 4,6E-02 |
| 1879 | superoxide dismutase [Fe] | -1,0355 | 4,7E-02 |
| 6144 | putative secreted protein | -1,0327 | 4,8E-02 |

**Table S 4:Strains used in this study, related to STAR Methods.**

| **name** | **description/ genotype** | **reference/source** |
|---|---|---|
| ***Escherichia coli* strains** | | |
| ***E. coli* NovaBlue** | general cloning host<br>*recA*1, *endA*1, *gyr*A96, *thi*-1, *hsd*R17, *supE*44, *relA*1, *lac* [F', proAB, lacIqZ, M15Tn10, (*tetR*)] | Novagen® |
| ***E. coli* ET12567//pUB 307** | host strain for conjugation from *E. coli* to *Kitasatospora* and *Streptomyces* (*dam*-13::Tn9, dcm-6, *hsdM*) ; pUB307, | [S1] |
| ***Kitasatospora purpeofusca* strains** | | |
| ***K. purpeofusca* ATCC 21470** | wildtype strain, producer of negamycin | ATCC strain collection |
| ***K. purpeofusca_∆neg1*** | *neg1* deletion mutant of *K. purpeofusca* | This study |
| ***K. purpeofusca_∆neg2part1*** | *neg2* deletion mutant of *K. purpeofusca* | This study |
| ***K. purpeofusca*_∆neg1::pRM4_*neg1.2*** | with *neg1.2* complemented *K. purpeofusca Δneg1* | This study |
| ***Streptomyces* strains** | | |
| ***S. lividans* TK24** | derivative of *S. lividans* 66 | [S2] |
| ***S. lividans*::pIJ_*neg1_ED::*BAC*neg2.2*** | heterologous host containing pIJ_*neg1 and* BAC*neg2.2* | This study |
| ***S. coelicolor* M1146** | derivative of *S. coelicolor* M145 | [S3] |
| ***S. coelicolor* ::pIJ_*neg1::BACneg2.2*** | heterologous host containing pIJ_*neg1 and* BAC*neg2.2* | This study |
| ***S. albidoflavus* Del14** | derivative of *S. albidoflavus* J1074 | [S4] |
| ***S. albidoflavus* ::pIJ_*neg1::BAC_neg2.2*** | heterologous host containing pIJ_*neg1* and BAC*neg2.2* | This study |
| ***S. albidoflavus*::pIJ_*neg1*** | heterologous host containing pIJ_*neg1* | This study |
| ***S. albidoflavus*::pIJ_ *BACneg2a*** | heterologous host containing BAC*neg2.2* | This study |

**Table S 5: Oligonucleotides used in this study, related to STAR methods.**

| no. | name | sequence | description |
|---|---|---|---|
| **1** | GAP_contigs_fw | ACCGCTCCGAGAACGAGGAGC | gap filling between the two resulting contigs of *K. purpeofusca* genome |
| **2** | GAP_contigs_rv | GTCTTCGACCCCGCCTACCT | |
| **3** | delneg1.1_up_fw | CTTCACATATGGTCGACTCGCTCACCACCTTC | deletion of *neg1.1* in *K. purpeofusca* |
| **4** | delneg1.1_up_rv | CCTATTAAGCTTGATTAGGGTCGCTGTGCTTTCAGGG | |
| **5** | delneg1.1_dw_fw | TAGGCTAAGCTTCTGGCGGCCGGCGAGAACGAG | |
| **6** | delneg1.1_dw_rv | GATTACGAATTCGTTCGACGGGTCCGTCGATCCG | |
| **7** | delneg1.1_contr.3_fw | GAACCGGTTCCGGCTGCGATG | control of *neg1.1* deletion in *K. purpeofusca* |
| **8** | delneg1.1_contr.3_rv | CCGACCAGGTGATGCTGCTG | |
| **9** | pRM4_compneg1.2_A_fw | GATCCCCCGGGCTGCAGGAATTCGATATCAAGCTTTCAGCCGGCGGGCGGGA | complementation of *neg1.1* in *K. purpeofusca* ∆*neg1.1* with genes 04339 - 04347 |
| **10** | pRM4_compneg1.2_A_rv | GCTAGCCAGGGGAGGACCCATATGGCCGTTCAGCTGAACCACACG | |
| **11** | pRM4_compneg1.2_B_fw | TGAACTTCAACATCCCGCCCGCCGGCTGAATGGAACTCACCTTCGTGGACGT | |
| **12** | pRM4_compneg1.2_B_rv | GGCCAGGTTGAGGAACGCCCCGCCGGCGAT | |
| **13** | pRM4_compneg1.2_C_fw | ATCGCCGGCGGGGCGTTCCTCAACCTGGCC | |
| **14** | pRM4_compneg1.2_C_rv | GATCCCCCGGGCTGCAGGAATTCGATATCAAGCTTCTAGTGGCGCAGCGCCCGGTC | |
| **15** | delneg2.1_up_fw | CATGCCATGGTACCCGGGAGCTCGAATTCGACACCAGGTGGTAGTAGGCGAGTTCG | deletion of *neg2.1* in *K. purpeofusca* |
| **16** | delneg2.1_up_rv | GGCCGGCCCGGTCGGCGGTGTGATGCTGCACTCCGCCCTCCAC | |
| **17** | delneg2.1_dw_fw | GGAGGGCGGAGTGCAGCATCACACCGCCGACCGGGCC | |
| **18** | delneg2.1_dw_rv | GAGCGGCCGCCACGGCGATATCGGATCCACTGGTGCTCGACGACTTCGGCACCG | |
| **19** | delneg2.1_contr._fw | CAGCCGCGCAGATTCTTCGCC | control of *neg2.1* deletion in *K. purpeofusca* |
| **20** | delneg2.1_contr._rv | GTCGGGATGCGAGTATTGCGGC | |
| **21** | pRM4_int_contr._fw | CTGGAAGTCCTCCATGGCCTGCTG | control genomic integration of pRM4 via ΦC31 attachement site |
| **22** | pRM4_int_contr._fw | GCGAGAAGCGCGACACGTCATAGAC | |
| **23** | CAPTURE_neg2.2_up1 | CGGTCGAAGGGCTTTGCCCGGGTGACGCGAGCACGACGAGACAGCGACACACTTGCATCG | for heterologous expression of *neg2.2* → cloning of BAC*neg2.2* using CAPTURE method |
| **24** | CAPTURE_neg2.2_dw2 | TCAACCAGTTCATTTGATCCGCAAGCCTTGACGAATGATGACGCTCAGTGGAACGAAAAC | |
| **25** | pBE45-universal-rv | ATCTTTATAGTCCTGTCGGGTTTCG | |
| **26** | pBE44/48-universal_fw | TTACCAATGCTTAATCAGTGAGGCACC | |

| | | | |
|---|---|---|---|
| **27** | gRNA_neg2.1_up1 | CCCGGGTGACGCGAGCACGAC | for heterologous expression of *neg2.2* → gRNA for BAC*neg2.2* cloning using CAPTURE method |
| **28** | gRNA_neg2.1_dw1 | ATCCGCAAGCCTTGACGAATG | |
| **29** | CAPTURE_BACneg2.2_test1_fw | GACGATCACGGTGTGGTTGCTG | for heterologous expression of *neg2.2* → proof of BAC*neg2.2* |
| **30** | CAPTURE_BACneg2.2_test1_rv | CGTGCTGGCCGTCAATCCG | |
| **31** | CAPTURE_BACneg2.2_test2_fw | CGGCCATGTTGAGACTGGCG | |
| **32** | CAPTURE_BACneg2.2_test2_rv | CAACTGGCCCTGTCGACGG | |
| **33** | CAPTURE_BACneg2.2_test3_fw | GACCTTCAGCCCGCGGTTC | |
| **34** | CAPTURE_BACneg2.2_test3_rv | GTTCAACGTCGAGTACCCGCAG | |
| **35** | pIJ_HetExp_creED_fw | GGTAGGATCGTCTAGAACAGGAGGCCCCATATGTGATCGACAGTCACCTCCAGATCTGTG | for heterologous expression of *creE*-like& *creD*-like |
| **36** | pIJ_HetExp_creED_rv | CATCTCGTTCTCCGCTCATGAGAACCTAGGATCCAAGCTTCTAGTGGCGCAGCGCCC | |
| **37** | hrdB_RNA_fw | CAAGCGCGAGCTGGAGATCATC | transcriptional analysis |
| **38** | hrdB_RNA_rv | GACCATGTGCACCGGGATACG | |
| **39** | 4344_RNA_fw | CGTTCCTGGTCACCGACGC | |
| **40** | 4344_RNA_rv | GGAGACATGGATGGCGATGGTCTC | |
| **41** | 4345_RNA_fw | CATCCAGATCGCGCCGAAC | |
| **42** | 4345_RNA_rv | GTCCTGCCGGTAGCCGG | |
| **43** | 4346_RNA_fw | GTCGGGACTGCCCTCGG | |
| **44** | 4346_RNA_rv | GTGCTGGAACGACTCTGCGC | |
| **45** | 4347_RNA_fw | GCTACCTGGAGTACGCCCGG | |
| **46** | 4347_RNA_rv | GTTGCGCTTGTGCGGCATC | |
| **47** | 8499_RNA_fw | AGTCCCAGTCGCTCCCGGAC | |
| **48** | 8499_RNA_rv | CTCGACGCCTGCGAAGCC | |
| **49** | 8500_RNA_fw | AAGCTCACCCATGCCCCGCC | |
| **50** | 8500_RNA_rv | GACCGTCACCGCACGCAG | |
| **51** | 8504_RNA_fw | CAGGTGACGGTGCCGCTGTC | |
| **52** | 8504_RNA_rv | GTCATCCTGGCAACCGGTTACC | |
| **53** | 2073_RNA_fw | GATCTGCTCACACCCGAGTTCTACG | |
| **54** | 2073_RNA_rv | CAGCACCTTGGTGGGGTAGC | |
| **55** | 8074_RNA_fw | GAAGCAGAAGCGGCAGTAGACC | |

**Table S 6: Plasmids used in this study, related to STAR methods.**

| name | Description | Reference/source |
|---|---|---|
| **pGusA21** | aac(3)IV, oriT, *ermEp** *gusA*, oripMB1A vector used for generation of gene deletion using blue white selection by β-glucoronidase | not published |
| **pGusA21_Δ*neg1.1*** | Deletion of the *neg1.1* region (4,726,499 bp-4,732,858 bp) in *K. purpeofusca* | this study |
| **pGusA21_Δ*neg2.1*** | Deletion of the *neg2.1* region (8,994,052 bp-8,999,551 bp) in *K. purpeofusca* | this study |
| **pRM4** | pSET152 derivative, *rep* (pMB1), *oriT*, *ΦC31 attP*, int, *ermEp**promoter, *aac*(3)IV Apra$^R$, expressions vector for streptomycetes | [S5] |
| **pRM4_*neg1.2*** | Genetic complementation of *neg1.2* (KPUR_ATCC21470_04339-04347) in *K. purpeofusca_Δneg1.1* | this study |
| **pIJ10257** | pMS81 derivative, *ermEp**, *ΦBT1attP*, *oriT* (from RK2), Hygro$^R$ | [S6] |
| **pIJ_*neg1*** | Heterologous expression of *creE*-like and *creD*-like genes. | this study |
| **pBE48** | Amp$^R$ | [S7] |
| **pBE45** | Apra$^R$ | [S7] |
| **BAC*neg2.2*** | BAC containing 27 kB region *neg2.2* including *neg2* used for heterologous expression | this study |

**Figure S1**

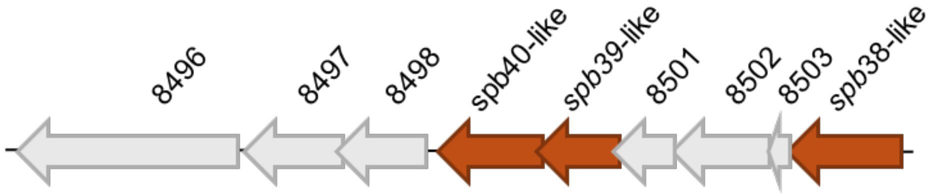

**Figure S 1: Candidate region 2 for negamycin biosynthesis (*spb* region).**
*spb* region encompassing *spb38-like* (KPATCC21470_8504) *spb39-like* (KPATCC21470_8500) and *spb40-like* (KPATCC21470_8501) homologous to *spb* genes involved in s56-p1 biosynthesis.

### Figure S2

A

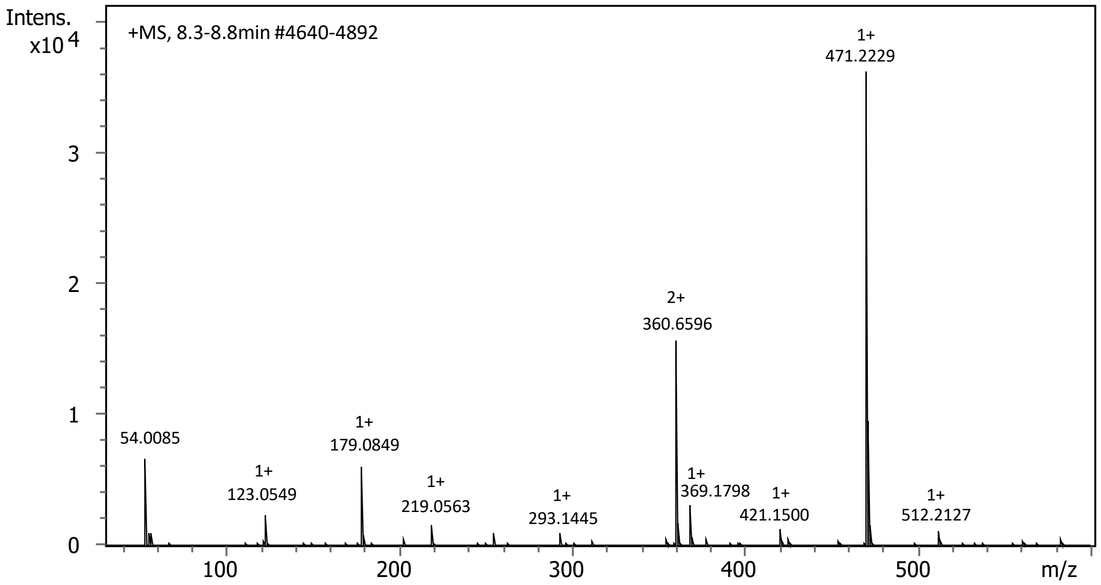
Intens.
x10 4
+MS, 8.3-8.8min #4640-4892
1+
471.2229
2+
360.6596
54.0085
1+
179.0849
1+
123.0549
1+
219.0563
1+
293.1445
1+
369.1798
1+
421.1500
1+
512.2127
m/z

B

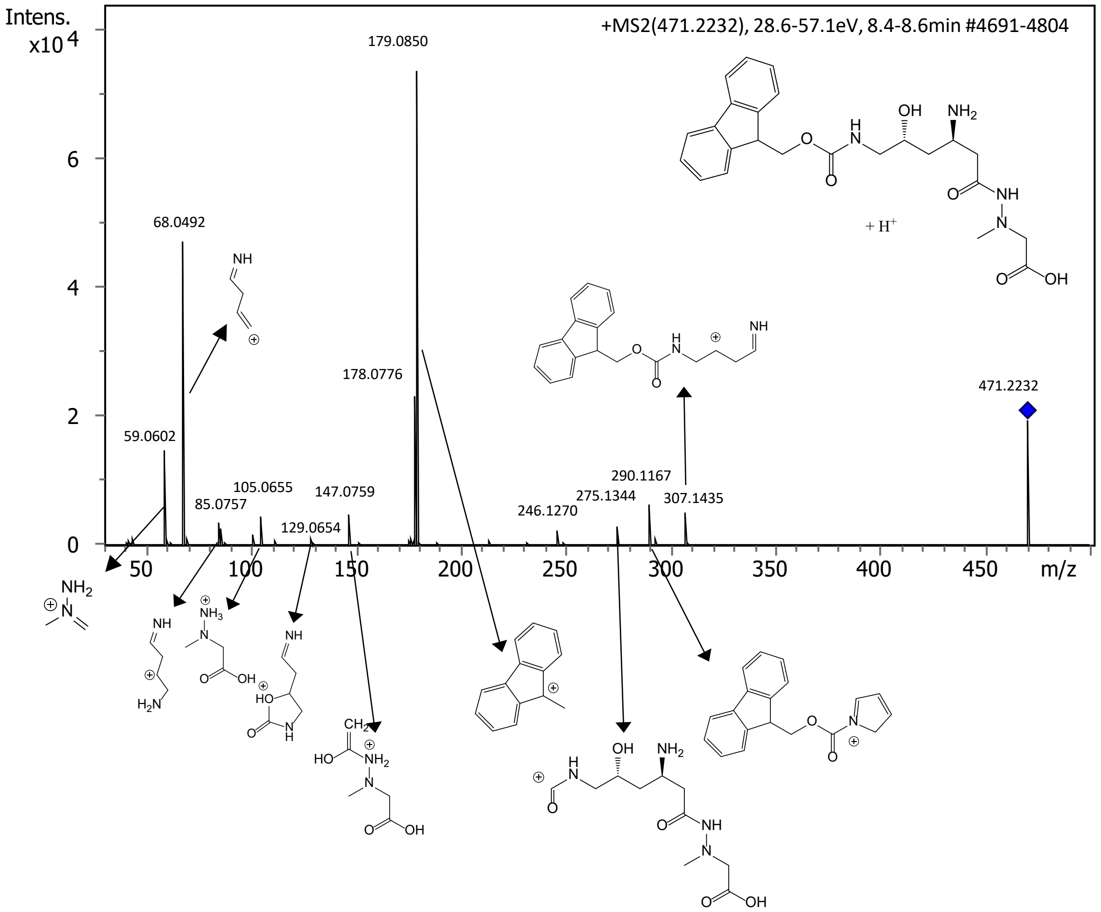
Intens.
x10 4
+MS2(471.2232), 28.6-57.1eV, 8.4-8.6min #4691-4804
179.0850
68.0492
178.0776
59.0602
105.0655
85.0757
147.0759
129.0654
246.1270
275.1344
290.1167
307.1435
471.2232
+ H+
m/z

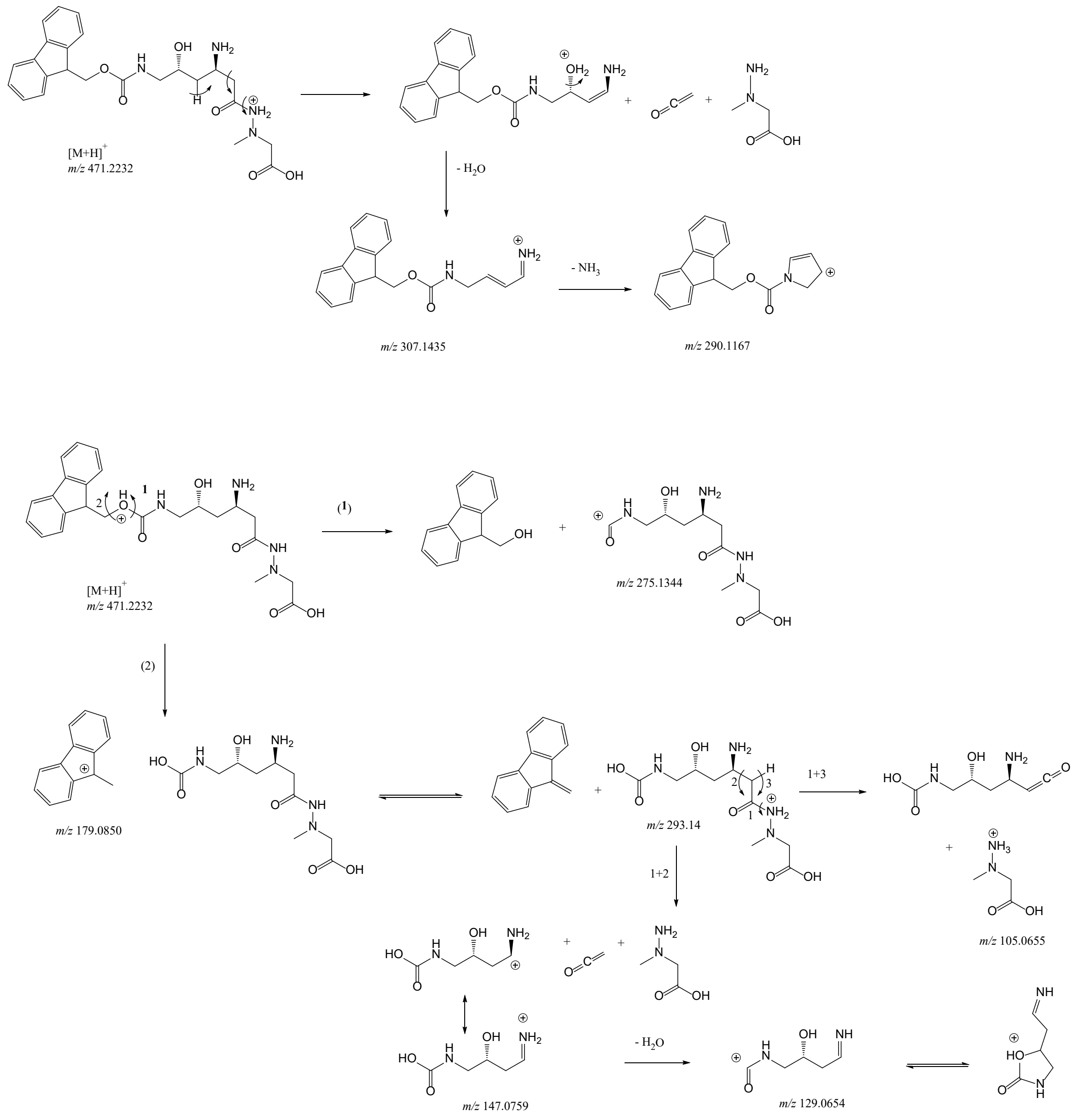

**Figure S 2:Mass spectrometry analysis of FMOC-negamycin, related to Figure 2.**
**A** HRMS spectrum of the peak detected at 8.5 min; **B** HRMS-MS of the ion at *m/z* 471.2229 $[M+H]^+$ annotated with the putative fragment ions, and **C** proposed fragmentation pathway of $[M+H]^+$

### Figure S3

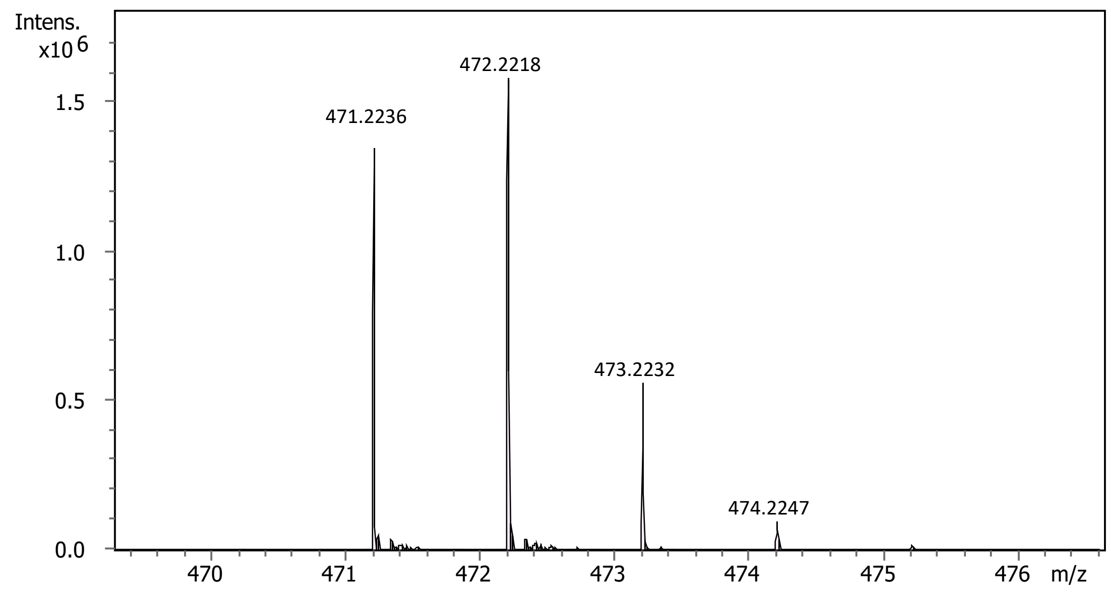

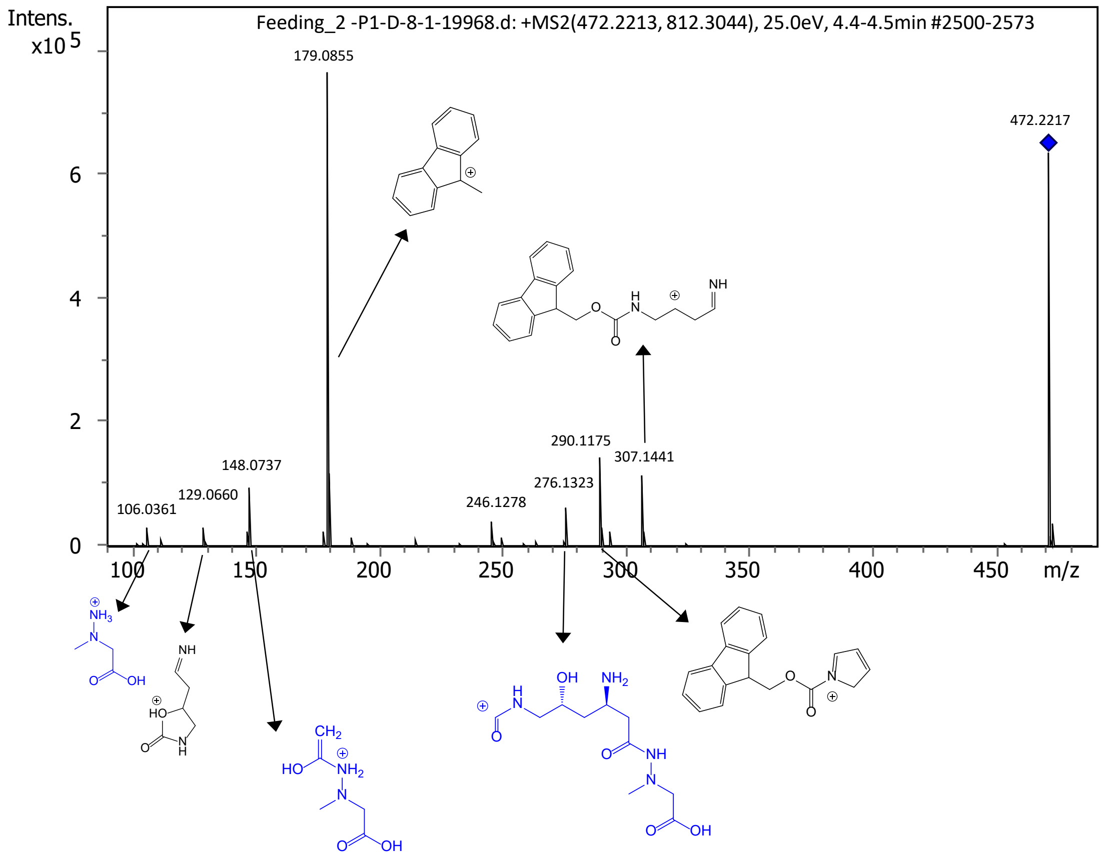

**Figure S 3: Incorporation of labeled $^{15}NO_2^-$ into negamycin.**
**A.** HRMS spectrum of FMOC-negamycin; **B** HRMS-MS spectrum of the ion at *m/z* $[M+H]^+$ 472.2218 annotated with the putative fragment ions.

### Figure S4

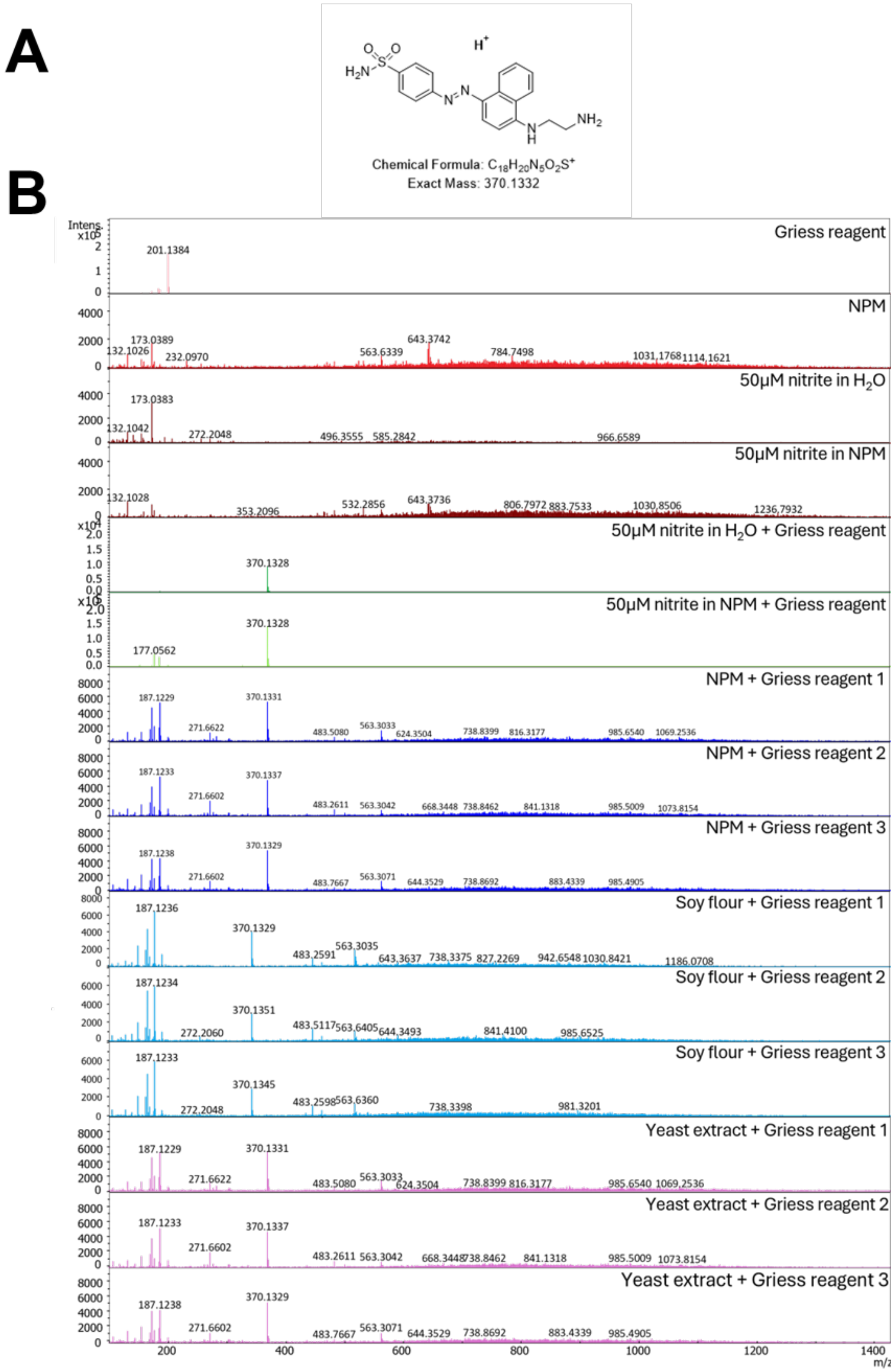

**Figure S 4:Nitrite detection in Negamycin Production Medium (NPM) using Griess reagent.**
A: Structure of azo product appearing if Griess reagent complexes with nitrite.
B: Mass spectra of HPLC-ESI-HRMS analysis; NPM and its components, yeast extract and soy flour, were tested for nitrite presence using Griess reagent. Samples were analyzed via HPLC-ESI-HRMS for the specific adduct (*m/z* $[M+H]^+$ 370.1329). As controls Griess reagent, NPM, NPM with added Griess reagent and water was analyzed. Each sample (NPM, soy flour and yeast extract) was analyzed in triplicates.

### Figure S5

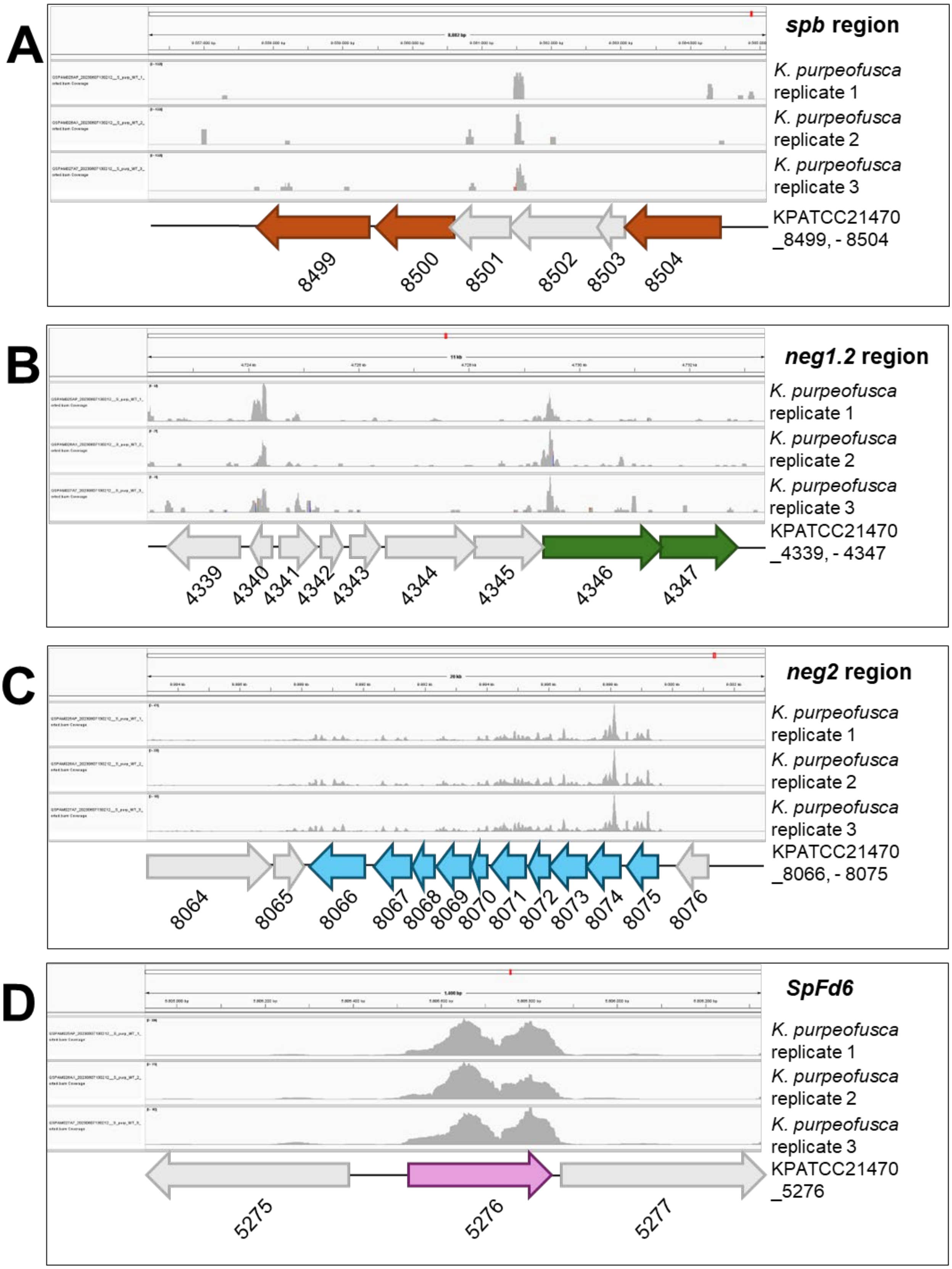
A
spb region
K. purpeofusca replicate 1
K. purpeofusca replicate 2
K. purpeofusca replicate 3
KPATCC21470 _8499, - 8504
8499
8500
8501
8502
8503
8504
B
neg1.2 region
K. purpeofusca replicate 1
K. purpeofusca replicate 2
K. purpeofusca replicate 3
KPATCC21470 _4339, - 4347
4339
4340
4341
4342
4343
4344
4345
4346
4347
C
neg2 region
K. purpeofusca replicate 1
K. purpeofusca replicate 2
K. purpeofusca replicate 3
KPATCC21470 _8066, - 8075
D
SpFd6
K. purpeofusca replicate 1
K. purpeofusca replicate 2
K. purpeofusca replicate 3
KPATCC21470 _5276
5275
5276
5277

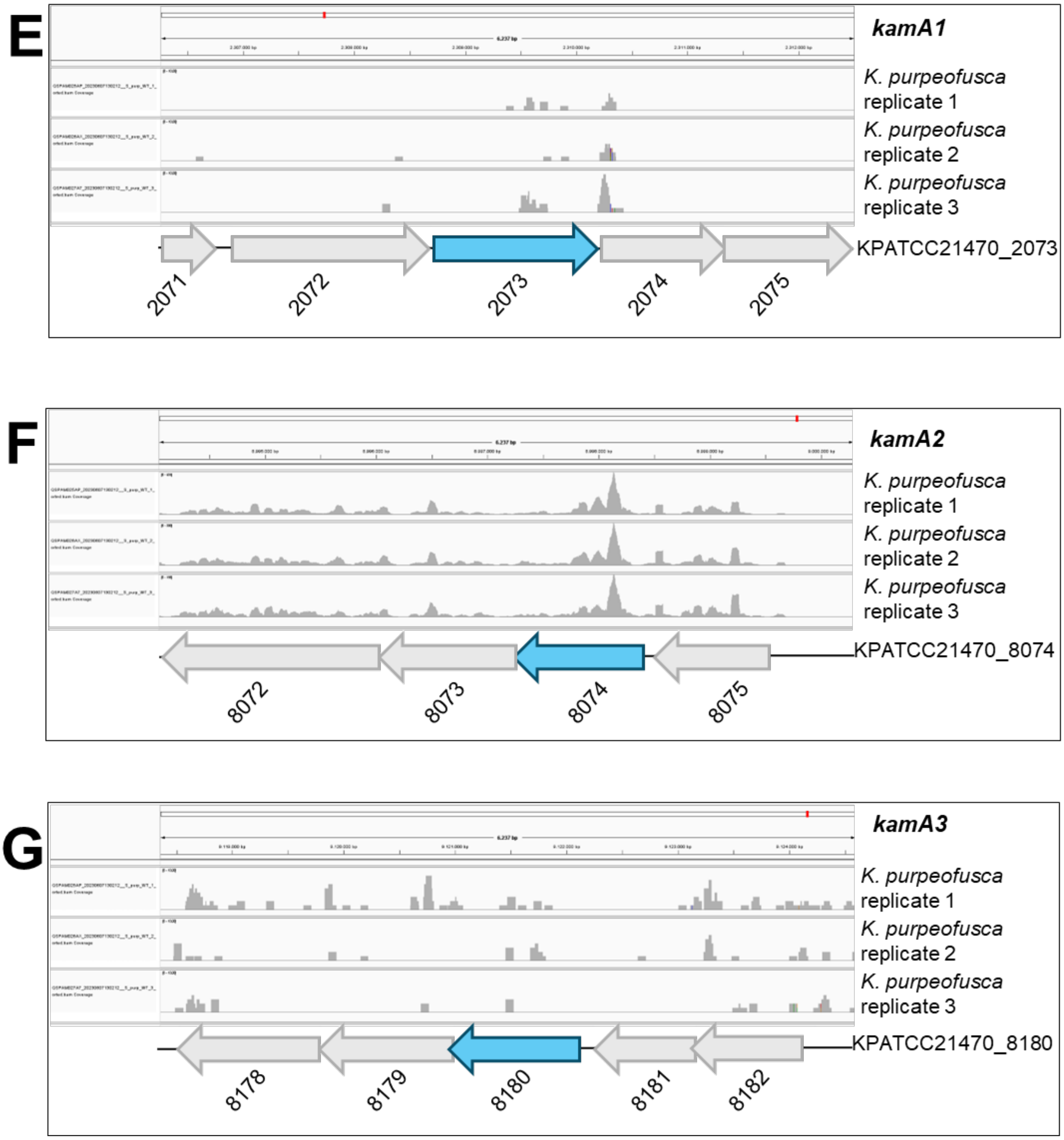

**Figure S 5: RNA-seq reads mapping in the studied regions (*spb*, *neg1*, *neg2* regions, *SpF6* and *kamA* homologs).**
RNA-seq reads are mapped against the genome sequence of *K. purpeofusca*.
A: Mapping of the reads in the *spb* region. *spb* homologs are highlighted in orange. B: Mapping of the reads in the *neg1.2* region. *creE* and *creD*-like genes are highlighted in green. C: Mapping of the reads in the *neg2* region. *neg2* genes are highlighted in blue. D: Mapping of the reads in the region encompassing *SpFd6* (purple). E: Mapping of the reads in the region encompassing *kamA1* (blue).F: Mapping of the reads in the region encompassing *kamA2* (blue). G: Mapping of the reads in the region encompassing *kamA3* (blue).

### Figure S6

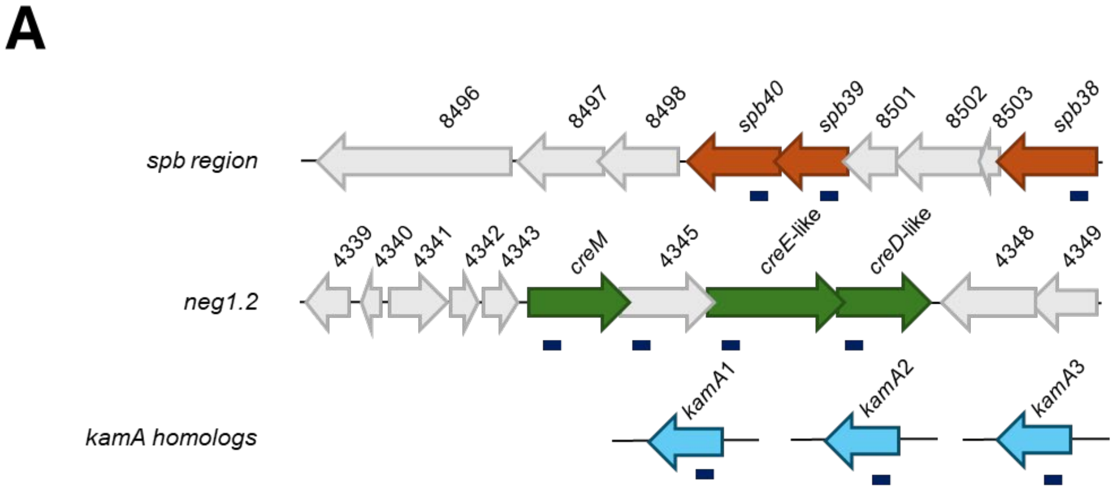

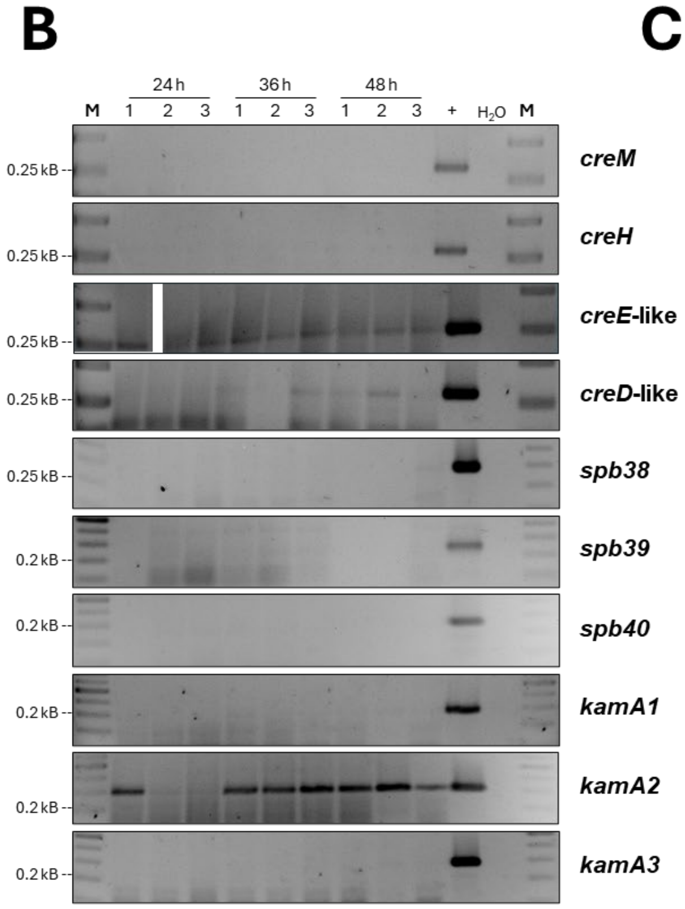

C

**Gene Transcription**

| | gene number | 24 h | 36 h | 48 h |
|---|---|---|---|---|
| *creM* | (4344) | no | no | no |
| *creH* | (4345) | no | no | no |
| *creE*-like | (4346) | yes | yes | yes |
| *creD*-like | (4347) | yes | yes | yes |
| *spb38* | (8504) | no | no | no |
| *spb39* | (8500) | no | no | no |
| *spb40* | (8499) | no | no | no |
| *kamA1* | (2073) | no | no | no |
| *kamA2* | (8074) | yes | yes | yes |
| *kamA3* | (8180) | no | no | no |

**Figure S 6:RT-PCR of *neg1.1* and *spb* regions and of *kamA* homologs.**
RT-PCR for amplification of 250-300 bp fragments was used to investigate the expression of the genes in *neg1* region (KPATCC21470_4344-4347), *spb* region (KPATCC21470_8504, KPATCC21470_8500 and KPATCC21470_8501) and of the *kamA* homologs *kamA1, kamA2* and *kamA3* (KPATCC21470_8074) after 24 h, 36 h 48 h growing under negamycin production conditions.
A: Fragments amplified are highlighted in blue.
B: Agarose gel pictures to visualize the amplification of fragments of the regions *in neg1.1* and *spb* region as well as *kamA* homologs. The images presented are cropped sections of the original gel to enhance the clarity and visualization of the results. In the case of the *creE*-like expression analysis, the second sample lane was omitted due to a pipetting error that affected the reliability of that particular result.
C: Expression of genes analyzed at time points 24 h, 36 h and 48 h.

**Figure S7**

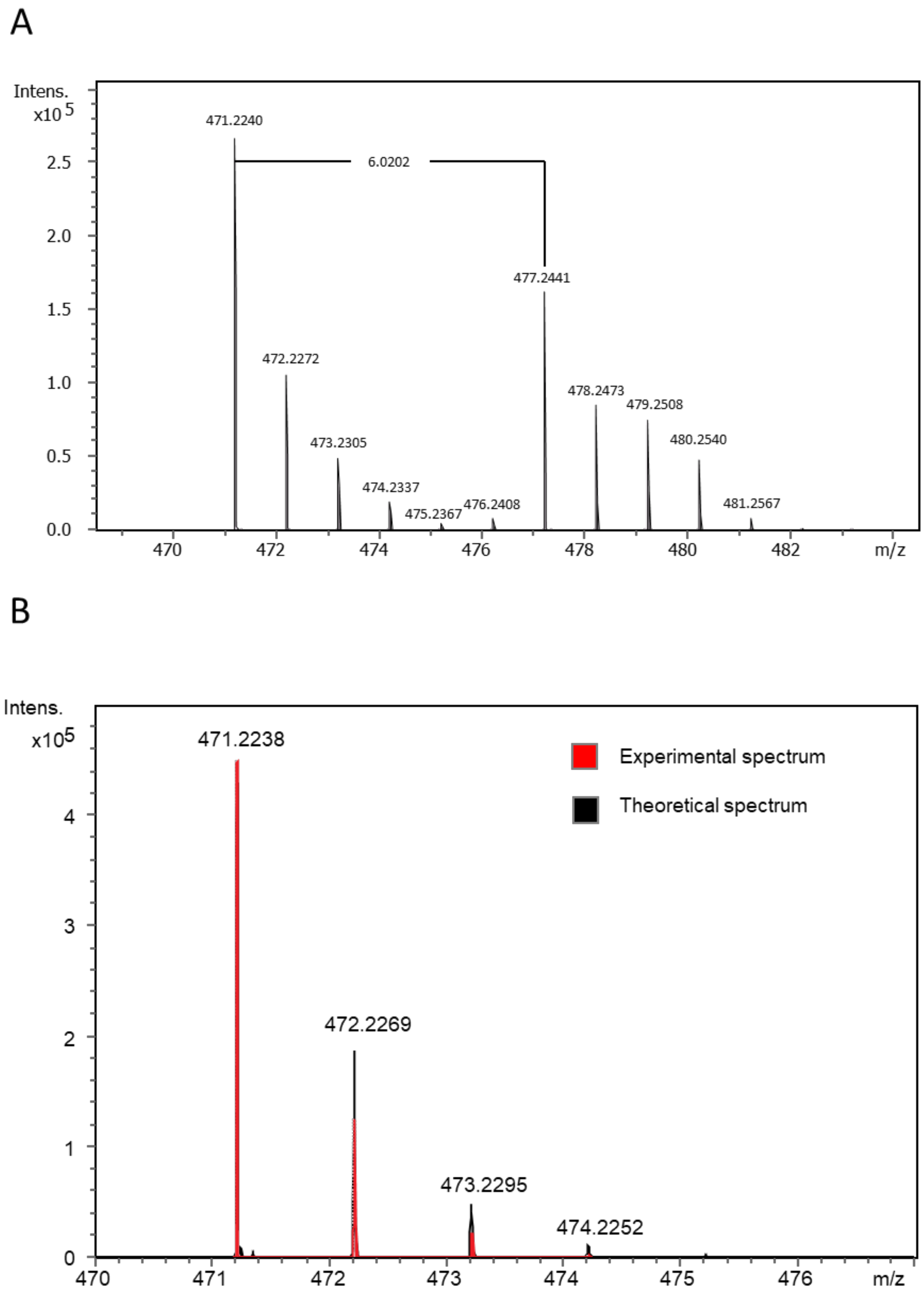
A
Intens.
x10 5
2.5
2.0
1.5
1.0
0.5
0.0
471.2240
6.0202
477.2441
472.2272
478.2473
479.2508
473.2305
480.2540
474.2337
475.2367
476.2408
481.2567
470
472
474
476
478
480
482
m/z
B
Intens.
x105
4
3
2
1
0
471.2238
Experimental spectrum
Theoretical spectrum
472.2269
473.2295
474.2252
470
471
472
473
474
475
476
m/z

C

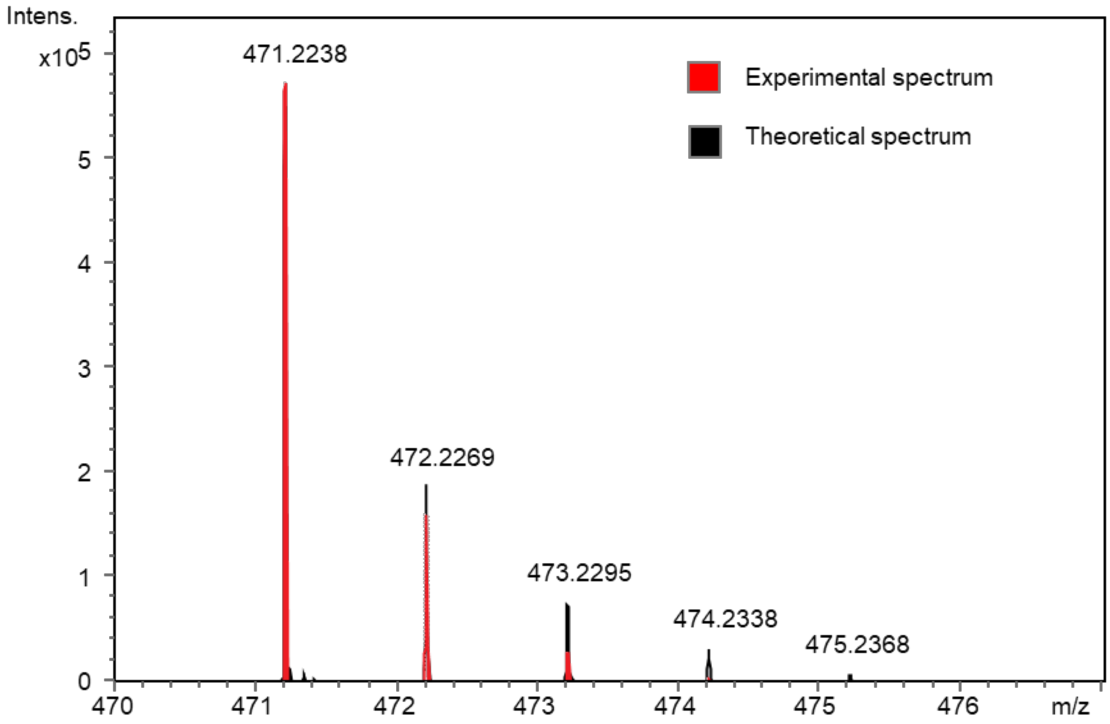

**Figure S 7:Incorporation of labeled $^{13}C_6$-L-lysine, $^{15}$N-glycine and $^{13}C_2$-glycine into negamycin.**
HRMS spectrum of FMOC-negamycin from treated culture supernatants of *K. purpeofusca* ATCC 21470 fed with **A**. $^{13}C_6$-L-Lysine, **B**. $^{15}$N-glycine and **C**. $^{13}C_2$-glycine.

### REFERENCES SI

[S1] Flett, F., Mersinias, V., and Smith, C.P. (1997). High efficiency intergeneric conjugal transfer of plasmid DNA from *Escherichia coli* to methyl DNA-restricting streptomycetes. FEMS Microbiol. Lett. 155, 223–229. https://doi.org/10.1111/j.1574-6968.1997.tb13882.x.

[S2] Rückert, C., Albersmeier, A., Busche, T., Jaenicke, S., Winkler, A., Friðjónsson, Ó.H., Hreggviðsson, G.Ó., Lambert, C., Badcock, D., Bernaerts, K., et al. (2015). Complete genome sequence of *Streptomyces lividans* TK24. J. Biotechnol. 199, 21–22. https://doi.org/10.1016/j.jbiotec.2015.02.004.

[S3] Gomez-Escribano, J.P., and Bibb, M.J. (2011). Engineering *Streptomyces coelicolor* for heterologous expression of secondary metabolite gene clusters. Microb. Biotechnol. 4, 207–215. https://doi.org/10.1111/j.1751-7915.2010.00219.x.

[S4] Myronovskyi, M., Rosenkränzer, B., Nadmid, S., Pujic, P., Normand, P., and Luzhetskyy, A. (2018). Generation of a cluster-free *Streptomyces albus* chassis strains for improved heterologous expression of secondary metabolite clusters. Metab. Eng. 49, 316–324. https://doi.org/10.1016/j.ymben.2018.09.004.

[S5]. Menges, R., Muth, G., Wohlleben, W., and Stegmann, E. (2007). The ABC transporter Tba of *Amycolatopsis balhimycina* is required for efficient export of the glycopeptide antibiotic balhimycin. Appl. Microbiol. Biotechnol. 77, 125–134. https://doi.org/10.1007/s00253-007-1139-x.

[S6] Hong, H.-J., Hutchings, M.I., Hill, L.M., and Buttner, M.J. (2005). The role of the novel Fem protein *VanK* in vancomycin resistance in *Streptomyces coelicolor**. J. Biol. Chem. 280, 13055–13061. https://doi.org/10.1074/jbc.M413801200.

[S7] Enghiad, B., Huang, C., Guo, F., Jiang, G., Wang, B., Tabatabaei, S.K., Martin, T.A., and Zhao, H. (2021). Cas12a-assisted precise targeted cloning using *in vivo* Cre-lox recombination. Nat. Commun. 12, 1171. https://doi.org/10.1038/s41467-021-21275-4.
